# False positive glycopeptide identification via in-FAIMS fragmentation

**DOI:** 10.1101/2023.05.28.542648

**Authors:** Valentina Rangel-Angarita, Keira E. Mahoney, Catherine Kwon, Raibat Sarker, Taryn M. Lucas, Stacy A. Malaker

## Abstract

High-field asymmetric waveform ion mobility spectrometry (FAIMS) separates glycopeptides in the gas phase prior to mass spectrometry (MS) analysis, thus offering the potential to analyze glycopeptides without prior enrichment. Several studies have demonstrated the ability of FAIMS to enhance glycopeptide detection but have primarily focused on N-glycosylation. Here, we evaluated FAIMS for O-glycoprotein and mucin-domain glycoprotein analysis using samples of varying complexity. We demonstrated that FAIMS was useful in increasingly complex samples, as it allowed for the identification of more glycosylated species. However, during our analyses, we observed a phenomenon called “in FAIMS fragmentation” (IFF) akin to in source fragmentation but occurring during FAIMS separation. FAIMS experiments showed a 2-5-fold increase in spectral matches from IFF compared to control experiments. These results were also replicated in previously published data, indicating that this is likely a systemic occurrence when using FAIMS. Our study highlights that although there are potential benefits to using FAIMS separation, caution must be exercised in data analysis because of prevalent IFF, which may limit its applicability in the broader field of O-glycoproteomics.

**Graphical abstract:** 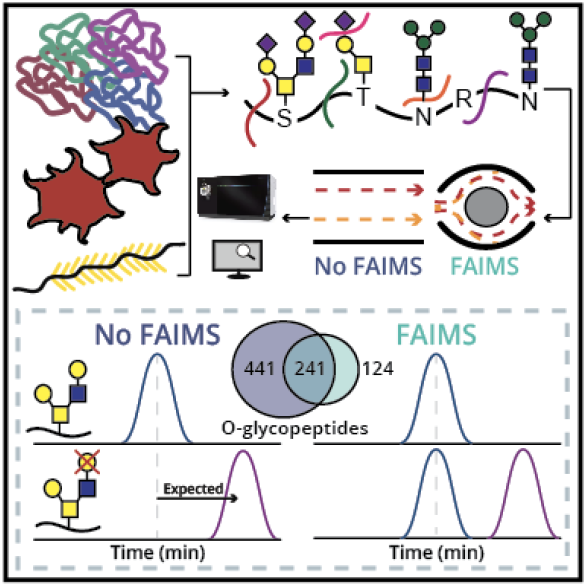

## Introduction

Post-translational modifications (PTMs) are used by eukaryotic cells to diversify their protein function and are important for different biological processes.^1^ Glycosylation is one of the most abundant PTMs, with over half of the human proteome predicted to be glycosylated.^2^ The two most common ways glycans modify proteins are through an N-linkage via an Asn residue or an O-linkage through the hydroxyl groups of Ser and Thr. N-glycosylation occurs within the consensus motif N-X-S/T, where X can be any amino acid except Pro. All eukaryotic N-glycans share a common chitobiose core structure (Man_3_GlcNAc_2_) and can be further extended and classified as high-mannose, complex, or hybrid.^3^ Mucin type O-glycosylation consists of an initiating N-acetylgalactosamine (GalNAc) and can be elaborated with other monosaccharides. As the name suggests, mucin-type O-glycosylation is prevalent within mucin domains, whose dense O-glycosylation forces the proteins to adopt a rigid bottlebrush-like structure with unique biophysical characteristics.^4,5^ Both N- and O-glycosylation play important biological roles. For instance, glycosylation is integral in viral infection,^6–8^ T cell activation and differentiation,^9^ and its dysregulation has been observed in a plethora of diseases.^4,10,11^

While mass spectrometry (MS) is the premier technique to characterize glycoproteins, the study of intact glycopeptides presents challenges not faced during typical proteomic studies. For instance, glycopeptides are often suppressed by unmodified peptides.^4,12^ Additionally, different glycoforms will decrease the signal observed of a given glycopeptide, an issue further compounded in glycopeptides with multiple glycosites.^12^ To this end, several enrichment and separation techniques can be exploited to enhance glycopeptide detection. Ion mobility spectrometry (IMS) is a widely used analytical technique that separates ions in the gas phase and has been shown to be useful in separating isomeric glycans, epimers, glycopeptides, and glycoprotein complexes.^13–18^ Here, analytes are passed through a region of high pressure drift gas under the effects of an electric field. Compact structures experience less collisions compared to extended structures, allowing for the resolution of mass, charge, and shape of analyte ions.^19^ High-field asymmetric ion mobility spectrometry (FAIMS) allows for fast gas-phase separation of ions between the electrospray emitter and inlet of the mass spectrometer.^20^ Here, an oscillating electric field is generated between two curved electrodes, which disperses ions based on mass, charge, shape, and dipole moment.^20^ A significant difference between the mobility of the species between high and low electric fields causes a net displacement of the ions towards the electrodes, resulting in neutralization. A compensation voltage (CV) affects the trajectory of different packets of ions, such that only a particular subset of species can enter the inlet of the instrument at a given time.^21^

Previous work showed that FAIMS allows for analysis of glycoproteins and glycopeptides that previously remained inaccesible.^20,22–24^ For example, Izaham *et al*. compared online FAIMS separation and offline zwitterionic hydrophilic interaction liquid chromatography (ZIC-HILIC) enrichment for bacterial glycoproteomes.^23^ Their results showed that while short aliphatic glycopeptides were not typically observed after HILIC enrichment, they could be readily identified using FAIMS.^23^ Ultimately, the combination of these two methods expanded the known glycoproteome of three *Burkholderia* species by >25%.^23^ Other studies showed FAIMS enhanced detection and confidence of glycopeptide assignments using tandem mass tag (TMT)-labeled N-glycopeptides,^24^ N-glycopeptides from alpha-1-acid glycoprotein,^25^ and both N- and O-glycopeptides from depleted human serum.^26^

As previous studies primarily concentrated on N-glycoproteomics, we targeted our analysis to O-glycoproteins and mucin-domain glycoproteins to determine if FAIMS could improve detection and separation of these often-overlooked species. In order to achieve this, we analyzed three samples consisting of a mucin-domain glycoprotein, a mixture of recombinant glycoproteins, and a platelet glyco-sheddome. FAIMS facilitated transmittance of precursors with higher charge states, which could enhance electron-based fragmentation, potentially leading to more localized O-glycosites. Our results showed that it was not particularly useful in the study of a single mucin-domain glycoprotein but did result in a higher number of glycopeptide identifications in the more complex samples. Notably, we observed fragmentation of glycosidic bonds during the FAIMS separation process, a phenomenon we termed “in-FAIMS fragmentation” (IFF). While this can happen during the typical electrospray ionization process (“in-source fragmentation”; ISF), we observed up to a 5-fold increase in the percentage of glycopeptide spectral matches originating from IFF relative to those from ISF. While FAIMS enabled identification of distinct glycopeptides, glycoforms, and glycoproteins in complex sample analyses, the presence of IFF raises concerns about the number of false positive identifications.

## Results and Discussion

### Podocalyxin

Mucin-domain glycoproteins are particularly challenging to study through MS techniques.^4^ Given their dense O-glycosylation and sequence composition, mucin domains are unamenable to digestion by traditional proteases.^4^ Therefore, these mucin domains are subjected to an additional digestion using mucinases, which cleave the protein backbone based on a combination of peptide sequence and glycan position.^4,27^ While this generates peptides from densely glycosylated stretches, they often have low charge density. This is caused by (a) absence of positive charge on the C-terminus, (b) large glycan structures, and/or (c) sequences mostly composed of amino acids with low proton affinity (e.g., Pro, Thr, Ser). To determine how these properties might affect FAIMS separation, recombinant podocalyxin was digested using two different mucinases: SmE^27^ and BT4244,^28^ both of which cleave N-terminally to glycosylated residues. While SmE can accommodate a variety of glycan structures in both the P1 and P1’ positions, BT4244 has a strong preference for desialylated core 1 structures.^27,28^ Therefore, the sample digested with BT4244 was co-treated with sialidase. As SmE digestion produced a wider variety of peptides and resultant identifications, the BT4244 results are primarily located in the supplemental information.

Our results focused on two features that can be used to determine the level of glycosylated species detected in a sample: glycopeptide spectral matches (GSMs), which represent the number of spectra identified by a search algorithm as glycosylated, and the number of unique glycopeptides (UGPs), which were generated by removing duplicates with identical peptide sequence and total glycan mass. With regard to SmE-digested podocalyxin, the number of identifications for each CV is displayed in **Figure 1A**, where the yellow bar graph represents UGPs and the overlapping line plot depicts GSMs. The experiments without FAIMS outperformed those using FAIMS at every CV tested for both GSMs and UGPs. That said, the CV setting that allowed for the most identifications was -50V, with 1290 GSMs and 269 UGPs, followed by -45V, with 1271 GSMs and 244 UGPs (**Table S1**).

**Figure 1.**
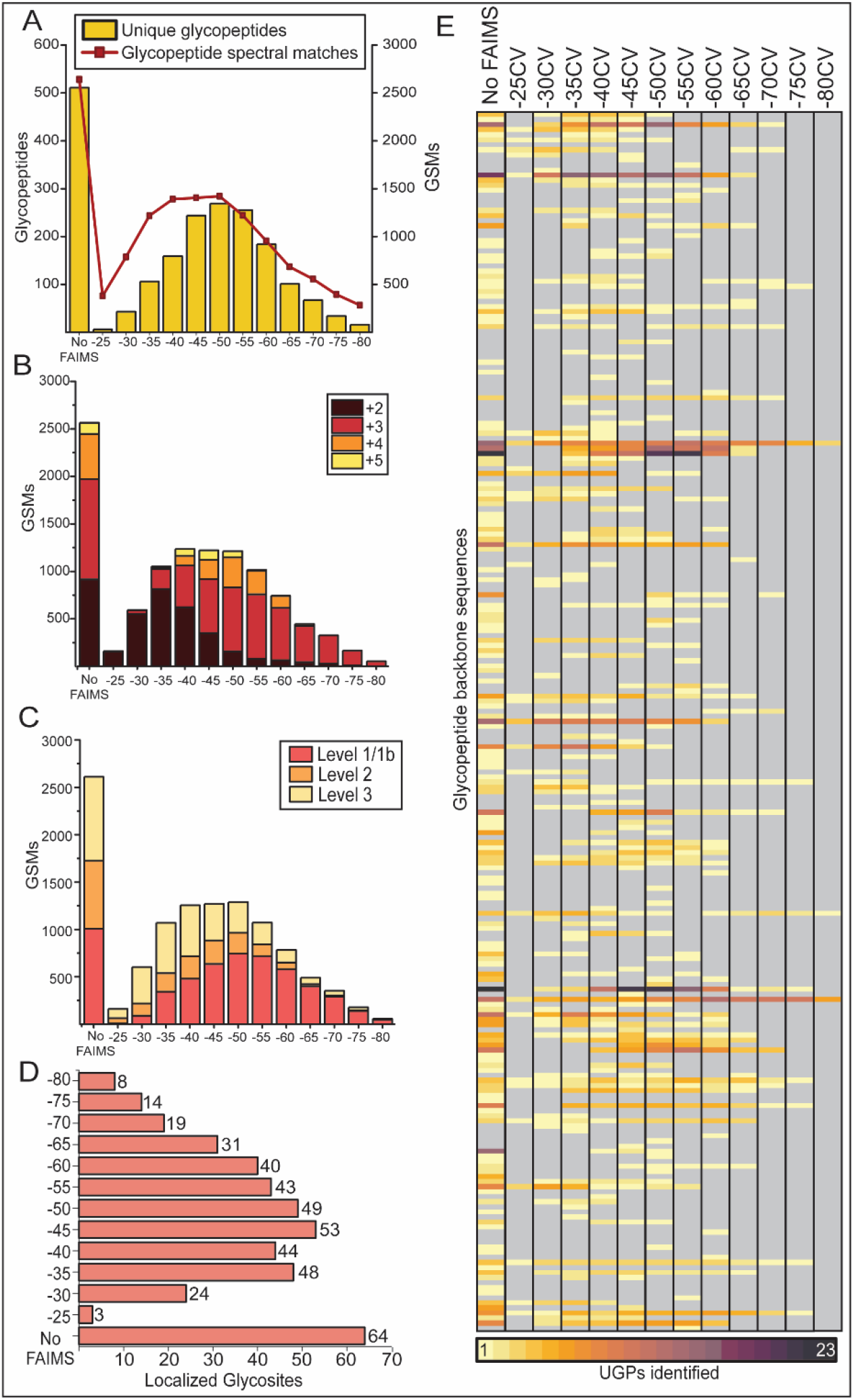
Benchmarking FAIMS separation using podocalyxin digested with SmE. Recombinant podocalyxin was digested with SmE and trypsin, then subjected to FAIMS separation and MS analysis. Resulting MS data was searched with O-Pair and manually validated. All figures were generated using OriginPro 2023. (A) GSMs (yellow, bar graph, right axis) and UGPs (red, line plot, left axis) identified with and without FAIMS. UGP lists were generated by removing duplicates of identical peptide sequences and glycan masses from search results. (B) Charge distribution of different FAIMS settings. GSMs were classified by precursor charge state, and results were plotted as a stacked bar graph. Identified charges range from +2 (dark red) to +5 (yellow). (C) GSMs were also classified based on the localization level assigned by O-Pair, where Level 1/1b (pink) described fully localized GSMs, while Level 2 (orange) represented those where at least one site could not be localized. Finally, Level 3 (yellow) illustrated GSMs where no glycan could be localized. (D) Following manual validation of GSMs, the number of localized glycosites per FAIMS setting was plotted. (E) Heatmap representation of UGPs identified for all peptide backbone sequences, where each row represented a unique glycopeptide sequence, and the color indicated the number of unique glycan masses attached to the peptide (legend at bottom). Gray boxes depicted glycopeptide backbone sequences that were not detected.

Electron-based dissociation (ExD) efficiency is highly dependent on charge density.^29^ An increase in precursor charge state of O-glycopeptides could yield better quality ExD spectra, thus allowing for improved glycan localization. Therefore, the precursor charge distribution, number of software-localized identifications, and validated glycosites were investigated. We hypothesized that an increase in CV magnitude (i.e., more negative) could lead to a proportional increase of overall glycopeptide charge, which we observed (**Figure 1B**). We then posited that the higher charge states could lead to improved spectral quality and thus better O-glycan site localization. To be sure, CVs more negative than -40V resulted in a higher proportion of localized identifications compared to the experiments without FAIMS (**Figure 1C**). A total of 70 O-glycosites were successfully validated and localized on podocalyxin; 64 were found without FAIMS analysis (91%). Using FAIMS, -45V yielded the most localized O-glycosites (53), though clearly fell short of the analysis without FAIMS (**Supplementary Dataset S1**). A small number of glycosites (6) were uniquely identified using FAIMS and were detected among the -45V and -50V CV datasets. These trends were replicated in the samples treated with BT4244 and sialidase (**Figure S1, Table S1, Supplementary Dataset S2** and **S3**). As demonstrated in **Figure 1E**, we generated a heatmap where each row represented a unique glycopeptide sequence and the color indicated the number of unique glycan masses attached to the peptide. Here, FAIMS settings allowed for the identification of up to 23 glycan masses (−45V and -50V), similar to those identified without FAIMS separation. Importantly, 73% (160) of the total glycopeptide backbone sequences (220) were identified without FAIMS.

Following an analysis performed by Izaham *et al*.,^23^ our results also showed that more negative CV values (−40V and lower) demonstrated a preference for longer peptides with a lower aliphatic index, while settings higher than -40V allowed for identification of shorter peptides with a higher aliphatic index (**Figure S2**). Interestingly, we found that FAIMS yielded peptides bearing smaller glycan compositions at all CVs measured (**Figure S2**). These trends were reproduced in podocalyxin digested with BT4244, albeit with less significance due to a lower number of identified glycopeptides (**Figure S3**).

Beyond its poor performance in glycopeptide analysis, we observed a significant decrease in signal when using FAIMS, which led us to inject larger sample amounts. To show this, five experiments at -45V were conducted using increasing amounts of sample ranging from 300 ng to 1200 ng. We then compared the extracted ion chromatograms (XICs) of the HexNAc oxonium ion for each experiment. The normalized intensity of the XIC for the 300 ng injection with FAIMS was 6% of the intensity observed without FAIMS (**Figure S4**). An increase in the injected amount led to an increase in the HexNAc XIC abundance, which correlated with a higher number of GSMs and UGPs, although this increase was less prevalent for settings beyond 725 ng (**Figure S5**). The sample amount can often be a limitation in glycoproteomic studies, especially when using clinically relevant samples. Therefore, the need for a larger injection amount could be restrictive when considering whether to use FAIMS. That being said, enrichment techniques like HILIC often require an even larger amount of starting material. For instance, Izaham *et al*. used 30 µg of protein for each ZIC-HILIC enrichment, compared to the 2 µg injected for FAIMS runs.^23^ As such, FAIMS may be preferable in lieu of performing an enrichment when sample amount is limited.

In summary, we showed that experiments without FAIMS allowed for the identification and localization of more glycosites when compared to those experiments using FAIMS. That said, in comparing FAIMS CV values, -45V and -50V generated the most UGPs and allowed for the localization of the most glycosites. Additionally, we observed a general increase of precursor charge state with concomitant increases in the proportion of localized glycosites by O-Pair; however, this increase in precursor charge state was not accompanied by an increase in the number of confirmed UGPs and glycosites. Given these observations, along with the increased sample and nitrogen requirements for FAIMS, we would not recommend routine use of FAIMS separation in single protein mucinomics workflows. If sample amounts allow, FAIMS could ostensibly be used to yield additional insight into these heavily O-glycosylated proteins, as demonstrated here.

### Recombinant Glycoprotein Mixture

To generate a sample more representative of those studied in the wider glycoproteomic field, we also analyzed a mixture of N- and O-glycopeptides from six glycoproteins. This sample had lower complexity when compared to the entire human proteome, which should allow more reliable search results while still providing a wider variety of peptide sequences.^30^ In accordance with previous CV optimization in other studies, ^20,21,23,25,26^ we performed single CV experiments from -25V to -80V in -5V intervals, as well as a control experiment without FAIMS. To select settings for double and triple CV experiments, we compared glycopeptides identified from single CV experiments, where we found that 87% of identifications were found using CVs -40, -45, or -50V. As such, a combination of these CVs were used for double and triple CV experiments (**Figure S6**).

Heat maps evaluating the number of UGPs showed that without FAIMS, 14 N- and 9 O-glycopeptide backbone sequences were identified, accounting for 63 and 54% of all N- and O-sequences observed. In the case of N-linked glycopeptides, 27% (5) of all detected glycopeptide backbone sequences (19) were found exclusively using FAIMS (−45V and -40/45V) (**Figure 2A**). Similarly, 44% (7) of all detected O-glycopeptide backbone sequences (16) were solely observed using FAIMS (−45V, -40/-50V) (**Figure 2B**). For both N- and O-glycopeptides, FAIMS identified more UGPs per glycopeptide backbone sequence, especially using CV settings of -40V and -45V, -40/-45V, and -40/-50V. As in the podocalyxin analysis above, we investigated the aliphatic indices of the peptides identified. While we observed some significance in the identification of shorter, less aliphatic peptides, a clear pattern was not observed (**Figure S7** and **S8**).

**Figure 2.**
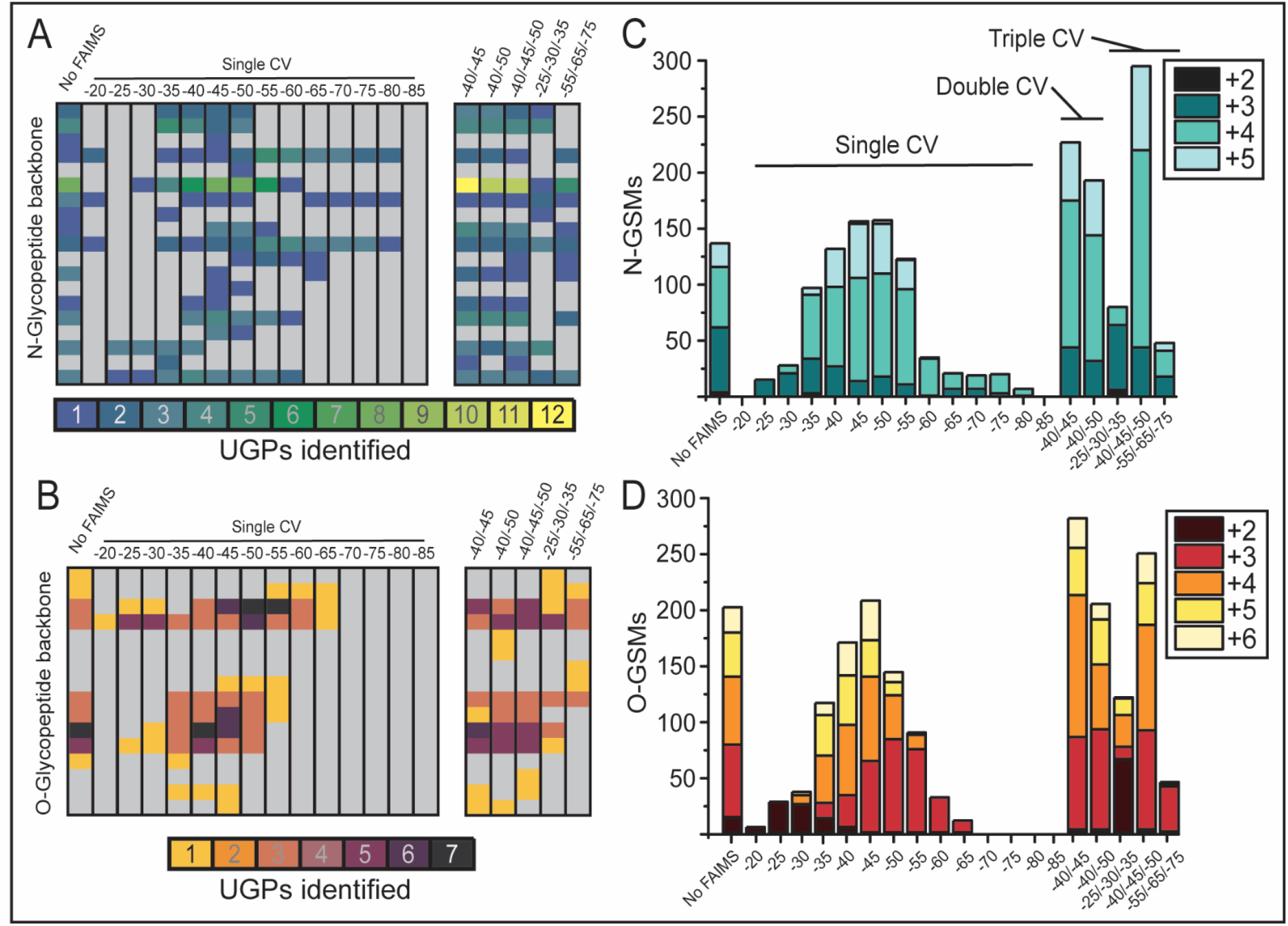
Glycoproteomics of a protein mixture using FAIMS separation. A mixture of N- and O-glycoproteins was digested with trypsin and subjected to FAIMS separation. Resulting MS data was searched with Byonic. All figures were generated using OriginPro 2023. (A) and (B) depicted heatmaps representing N- and O- UGM per peptide, respectively, where each row represented a unique glycopeptide sequence, and the color pictured the number of UGMs attached to the peptide (legend at bottom). Gray boxes depicted glycopeptide backbone sequences that were not detected when using a given CV. (C) and (D) depicted N- and O-GSMs identified across different FAIMS settings, respectively, as well as the charge state distribution for each CV. Charge state distribution is depicted as a stacked bar graph, where the darkest color portrayed the lowest charge observed (+2), and lighter represented higher charges (light blue depicts +5 for C, light yellow depicts +6 for D).

In contrast with our results above, we noted a rise in the number of GSMs identified using FAIMS experiments, though the peptide charge states also exhibited a similar upward trend (**Figure 2C** and **2D, S9**). Using -45V yielded 164 N-GSMs and 203 O-GSMs, which represented a 6% and 7% increase, respectively, compared to experiments without using FAIMS. The setting of -50V generated the most N-GSMs (166, 8% increase); however, we were only able to identify 141 O-GSMs, indicating that -45V may be preferable to enhance O-glycopeptide detection. Employing double and triple CV settings enabled an even larger difference in the quantity of GSMs. For example, -40/-45V demonstrated a 47% increase in N-GSMs (227) and a 50% increase in O-GSMs (284) compared to experiments without FAIMS. Finally, the triple CV setting of -40/-45/-50V allowed for the identification of 92% more N-GSMs (295) and 33% more O-GSMs (252). Despite the enhanced number of glycosylated spectra, only -40/-45V resulted in higher N-UGPs when compared to experiments without FAIMS (44 N-UGPs, 13% increase). Similarly, -45V was the only setting where O-UGPs were higher (28 O-UGPs, 12% increase) (**Figure S9, Table S3**, and **Table S4**). Thus, the augmented number of GSMs was likely caused by redundant glycopeptide spectra.

Ultimately, when comparing FAIMS to control experiments, one-third of glycopeptide backbone sequences were identified exclusively using FAIMS separation. We also observed more UGPs per backbone sequence. Based on our results, we recommend a single CV of -45V for O-glycoproteomic analysis, and -50V for N-glycoproteomics. Importantly, while double and triple CVs yielded a higher number of GSMs, they suffered in identification of UGPs compared to single CV experiments due to redundant GSMs. Nevertheless, FAIMS separation permitted a better coverage of the glycoproteome of this sample compared to experiments without FAIMS, demonstrating its potential utility in glycoproteomic workflows for a mixture of proteins.

### Glycoprotein enrichment of the platelet sheddome

To explore the application of FAIMS to a highly complex sample, we took advantage of lectin enriched platelet sheddome isolated from human blood,^31^ which presented a more intricate blend of N- and O-glycosylation. These experiments used the settings that yielded the highest number of identifications in the first two sample types (−40V, -45V, -40V/-45V, and -40V/-50V). The use of FAIMS significantly improved the number of GSMs, UGPs, and glycoproteins identified for both N- and O-glycans. FAIMS at -40V allowed for the detection of 449 GSMs and 126 UGPs, as opposed to the 196 GSMs and 85 UGPs identified without FAIMS (**Figure 3A**). The majority of glycopeptide backbone sequences were detected in experiments conducted without FAIMS, but many could be accounted for using -40V (93% of O- and 89% of N-glycopeptides, **Figure 3B**). No additional proteins were identified outside of CV -40V and the experiment without FAIMS (**Figure 3C**). Overall, the addition of FAIMS yielded complementary results to those found without it, significantly increasing the glycoproteomic coverage of this sample.

**Figure 3.**
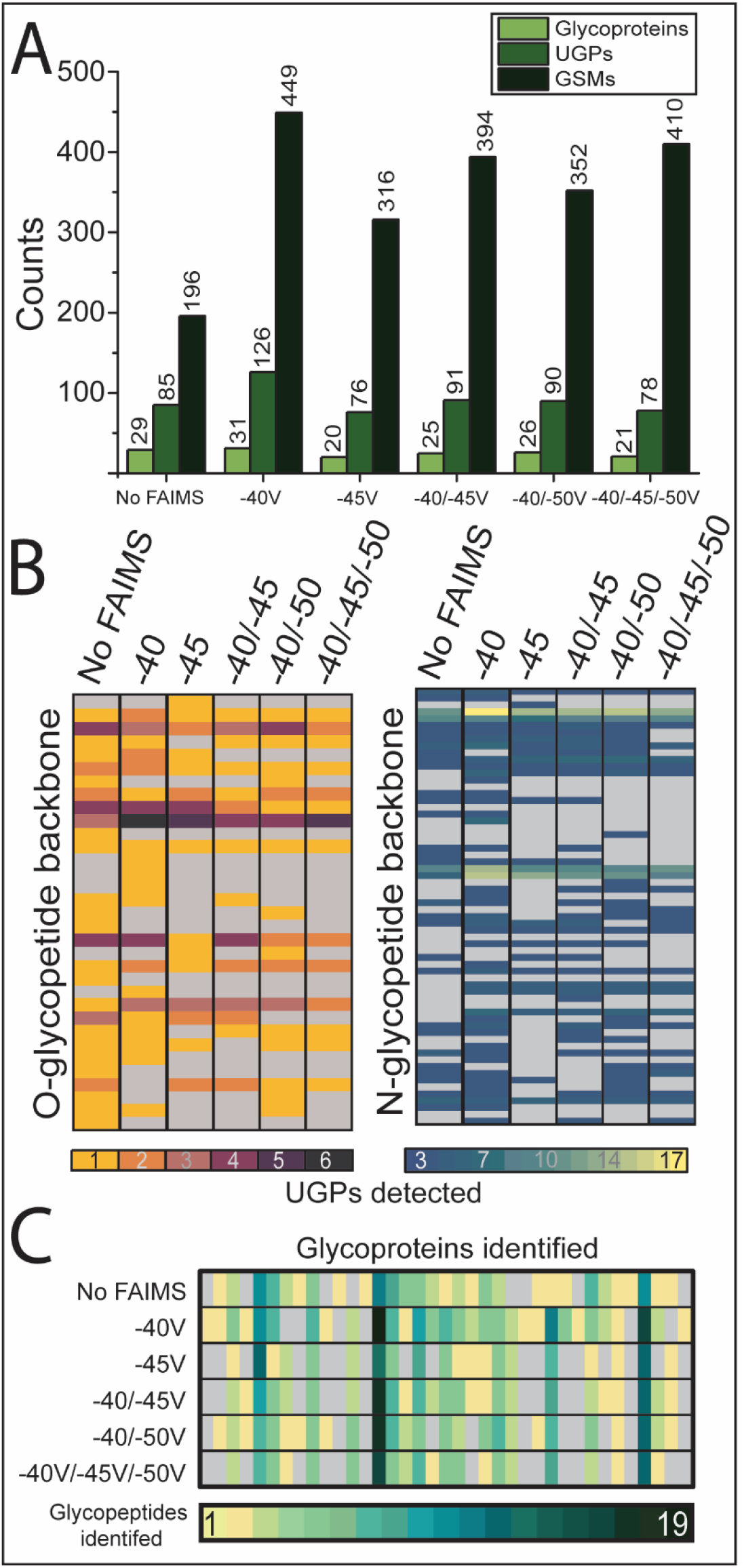
Glycoproteomic analysis of FAIMS separation in a highly complex sample. Experiments with and without FAIMS were performed in a sample containing a lectin enriched platelet glyco-sheddome. Following data acquisition, MS data was searched using Byonic and results were filtered to ensure confident glycopeptide backbone identification. (A) Depicted glycoproteins (light green), UGPs (green) and GSMs (darker green) identified with a range of FAIMS CVs and without FAIMS. (B) Heatmaps showing O- (left) and N- (right) glycoprotein backbone sequences, where the z axis pictured the number of glycoforms identified. (C) Heat map of all glycoproteins identified in each CV setting, where each column represented an individual protein and each row described a different CV setting. The colors illustrated the number of UGPs identified for each protein, with a legend shown below.

Interestingly, we noted that performing double and triple CV experiments often reduced glycoproteomic coverage at the protein level compared to the single CVs. In these experiments, the total cycle time is equal to that of a single CV at 3 seconds. Therefore, MS2 spectra for each CV were collected for a shorter period of time, which could have resulted in only the most abundant species being selected for fragmentation. Since dynamic exclusion is applied to each CV separately, the same species could be selected in multiple CVs during a double or triple CV experiment, yielding more GSMs but fewer UGPs.

Finally, we observed in initial experiments that the chromatogram without FAIMS was dominated by polyethylene glycol (PEG). Since PEG elutes on the same chromatographic timescale as our peptides of interest, its abundance and ionizability can cause suppression of peptide signal, particularly for lower abundance and/or poorly ionizing species such as glycopeptides. We selected CVs -35V, -45V, and -55V to study the ability of FAIMS to reduce the presence of PEG observed during MS analysis. Adding a CV of -35V led to lower PEG signal which continued to decrease alongside the CV, as shown in **Figure S10**. While all FAIMS settings yielded more GSMs (up to 316 GSMs) compared to the control experiment (196 GSMs), only -35V demonstrated a slight increase in UGPs (86 UGPs) relative to the experiment without FAIMS (85 UGPs). While FAIMS reduced apparent polymer contamination in the gas phase, performing FAIMS fractionation after HPLC separation means that any contaminants present during chromatographic separation could potentially affect both the elution of peptides and the longevity of the columns used. Additionally, such contaminants may still suppress the ionization of glycopeptides during the electrospray process.

### In-FAIMS Fragmentation

Glycosidic bonds are very labile in nature, as evidenced by the fact that glycopeptide spectra generated using collisional fragmentation methods are often dominated by glycan losses from the precursor ion.^32^ As a result of this, and the presence of drift gas during FAIMS separation, we suspected that IFF could be occurring in our analyses. These thoughts were corroborated by: (a) observations of fragmentation during FAIMS analysis of glucose and levoglucosan^33^ and (b) possible ion heating.^34–38^ Briefly, under high electric fields, the ion’s drift velocity exceeds its thermal velocity, which will lead to inelastic collisions that convert translational energy into internal energy. As a consequence, fragmentation within the IMS cell can occur, resembling collisional dissociation methods.^34–38^ Ion heating can cause structural changes and fragmentation, and has been extensively studied in other IMS techniques. ^34–38^

We assessed the XICs of the ten most abundant glycopeptides in podocalyxin acquired with -45V, as well as the masses corresponding to glycan losses from the initial glycosylated precursor ion. Further, we compared abundance of these glycopeptides in runs with and without FAIMS; note that the latter would simply be observations of ISF. For this analysis, we considered positive instances of source fragmentation (ISF or IFF) if the chromatographic profile of the complete precursor ion and the glycan losses were identical. This suggested that peptides with different glycan compositions were eluting at the same time and with a similar pattern, a trend unlikely to occur without ISF/IFF. To confirm IFF or ISF, we used the XIC of masses corresponding to potential false-positive glycopeptides, alongside those from glycopeptides with equal peptide backbone and larger glycan masses. We observed their elution profiles relative to peptides with larger glycans and the same peptide sequence, and if their chromatographic profiles were identical, these were noted as ISF/IFF.

As an example, **Figure 4A** and **4B** depicts two instances of IFF where, in this case, the loss of a sialic acid (NeuAc, A) is observed. Glycopeptides without sialic acid are expected to elute earlier in the gradient, given the decrease in their hydrophilicity. In both examples, the chromatograms display two peaks for the precursor with a smaller glycan mass: one corresponding to the glycopeptide not resulting from IFF, which elutes earlier in the gradient. The other peak is the product of IFF, as evidenced by the similar chromatographic profile to the glycopeptide with one more sialic acid. While it is possible that different glycoforms of one glycopeptide elute at similar retention times, they can typically be distinguished by their chromatographic profile, and therefore are not annotated as ISF/IFF (**Figure 4A**).

**Figure 4.**
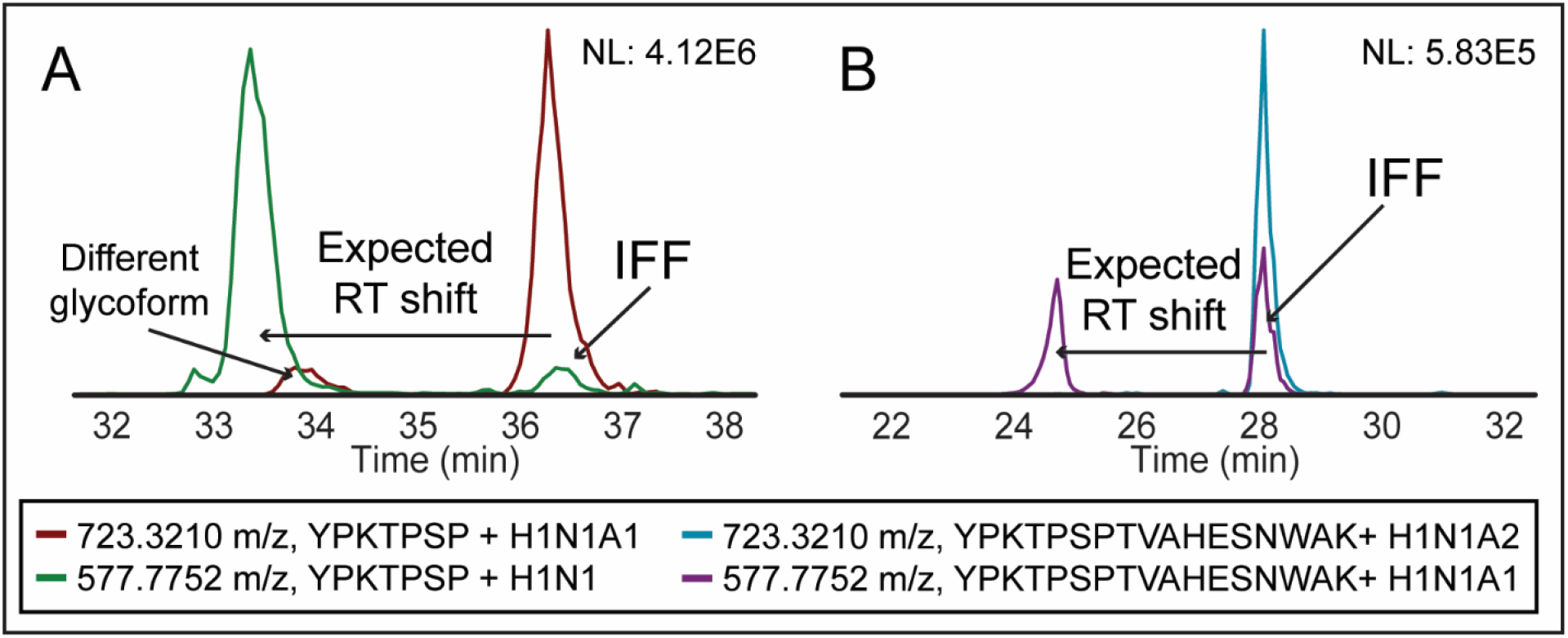
Assignment of ions product of source fragmentation. Recombinant podocalyxin was digested with mucinase SmE and trypsin. (A) and (B) depicted chromatograms with IFF, both acquired with a CV of -45V. (A) The peptide YPKTPSP with glycan masses HexNAc1-Hex1-Sia1 (red) and HexNAc1-Hex1 (green). (B) The peptide YPKTPSPTVAHESNWAK with glycan masses HexNAc1-Hex1-Sia2 (blue) and HexNAc1-Hex1-Sia1 (purple). Both figures were annotated with expected retention time shift without ISF/IFF (left) and evidence of IFF (right).

Out of the ten most abundant glycosylated precursors, we observed evidence for IFF in two instances, including losses of hexose (Hex, H) and N-acetylhexosamine (HexNAc, N). For example, ISF and IFF of the peptide SSQMPA with glycan HexNAc2-Hex2 (675.7711 m/z, +2) was observed with glycan masses corresponding to HexNAc2-Hex1, HexNAc1-Hex1, and HexNAc2 (**Figure 5A** and **5B**). As depicted by the chromatographic trace in blue, we observed the loss of hexose (blue) in both experiments, but the intensity of this loss was much lower without FAIMS (**Figure 5A**, 8.5% of precursor intensity), when compared to the experiment with FAIMS (**Figure 5B**, 60.5% of precursor intensity). Further, the chromatographic peak corresponding to the loss of two hexoses was present in FAIMS (pink) at an abundance of 57.8% relative to the precursor ion (**Figure 5B**); this mass was negligible in the experiment without FAIMS (**Figure 5A**). Importantly, the intact glycopeptide abundance was significantly lower in the FAIMS experiment, likely due to the presence of IFF. Similar results were found on peptide TSPATALR with glycan HexNAc2-Hex2 (773.8618 m/z, +2), as it was observed in the same chromatographic window with glycan masses corresponding to HexNAc2-Hex1, HexNAc2, and HexNAc1 (**Figure S11**).

**Figure 5.**
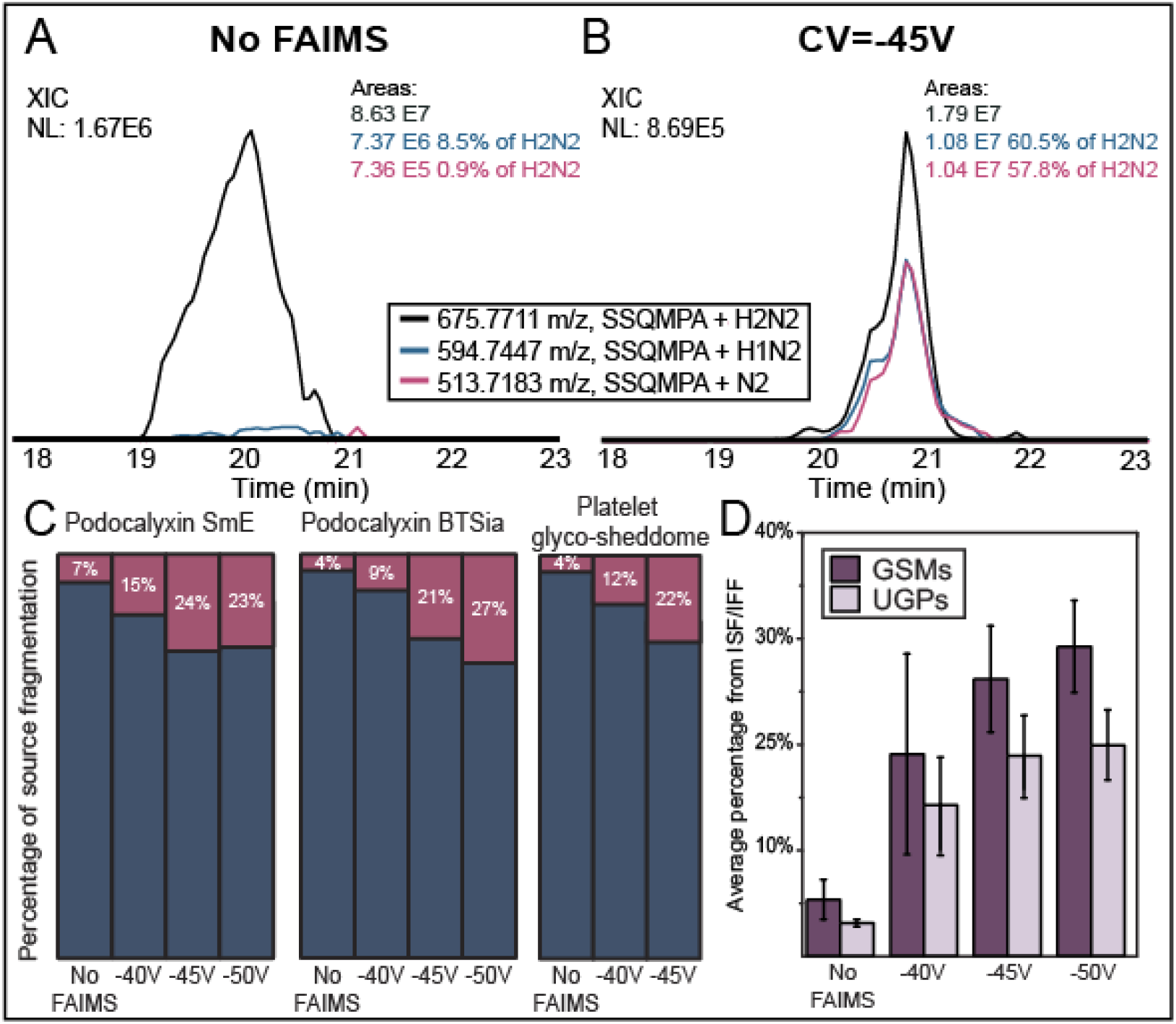
FAIMS experiments exhibit a larger number of identifications from IFF. Recombinant podocalyxin was digested with SmE and trypsin. Using Thermo XCalibur, we extracted ion chromatograms of the doubly charged glycopeptide with glycan masses HexNAc2-Hex2 (black), HexNAc2-Hex1 (blue), and HexNAc2 (pink) on peptide SSQMPA. The abundance percentage of each peak was calculated relative to that of the original precursor species. (A) Extracted chromatograms from the experiment without FAIMS and (B) those acquired using CV -45V. Alongside each chromatogram, the glycopeptide and glycan composition, m/z, integrated area, and normalized abundance of peaks assigned as IFF are shown. (C) Normalized percentage of GSMs (pink) caused by ISF/IFF. All spectral matches that eluted within a minute of another identification with the same backbone and larger glycan mass were evaluated to verify if they originated from ISF/IFF. Following, the percentage of these instances of ISF/IFF was calculated relative to the total number of identifications. This analysis was performed in data acquired under control conditions, and single CVs -40V, -45V and -50V. The samples analyzed were podocalyxin digested with SmE (left) and BT4244 (middle) and the lectin enriched platelet sheddome (right). (D) Average GSMs (dark purple) and UGPs (light purple) that originated from ISF/IFF from all samples analyzed. Standard deviation is depicted using error bars.

To establish this behavior on a larger scale, we developed “Glyco-SourceFragFinder”, a program that detects possible occurrences of both IFF and ISF by marking identifications that elute within a minute of a peptide with the same backbone sequence but with larger glycan masses. To manually inspect the resulting lists, we extracted the XICs of masses corresponding to the potential IFF/ISF glycopeptides and the precursor ions. IFF/ISF was assigned if the chromatographic profiles were the same.

We confirmed a larger proportion of total identifications (GSMs and UGPs) originated from IFF in FAIMS experiments compared to those generated from ISF in the control experiment (**Supplementary Dataset S7**). For instance, in the SmE digest of podocalyxin, 188 GSMs were confirmed as ISF without FAIMS, whereas FAIMS experiments yielded up to 283 validated IFF GSMs (**Table S5, Figure S12A**). ISF accounted for only 7.3% of all GSMs without FAIMS, while IFF resulted in a significant increase in false positive identifications (15.4% for -40V, 24.1% for -45V, and 23.3% for -50V, **Figure 5C**, pink). A similar trend was observed in UGPs, where 6% were caused by ISF for experiments without FAIMS, compared to up to 20.1% detected at -50V via IFF (**Figure S12A**). These results were also replicated in podocalyxin digested with BT4244 (**Table S6, Figure 5C**, pink, **Figure S12B**) and the platelet glyco-sheddome (**Table S7, Figure 5C**, pink, **Figure S12C**). The corrected GSM counts (i.e., true positive identifications) are depicted in **Figure S12D**.

To investigate whether these observations were unique to our instrumentation setup, we repeated the above analyses with data from depleted human serum used in a recent preprint by Alagesan *et al*.^26^ MS files were downloaded from PRIDE (identifier PXD038673), searched with Byonic, and analyzed in the manner described above. Our results demonstrated that on average, 39.7% of GSMs originated from IFF in -40V, 44.6% in -45V, and 47.3% in -50V (**Table S8, Figure S13**). These values were consistent across two replicates; using -45V, 83 GSMs were the product of IFF in the first replicate and 82 in the second. Notably, the observed increase in IFF relative to our data could have been caused by the electrode temperature. While our experiments were performed with an inner electrode temperature of 70 °C, data collected by Alagesan *et al*. ^26^ was acquired at 100 °C. The higher temperature could increase ion heating of glycopeptides as they are separated in the FAIMS source, thus leading to more instances of IFF.

Overall, we observed a two- to five-fold increase in glycan fragmentation when using FAIMS separation (**Table S9, Figure 5D**) when compared to experiments without FAIMS, which was replicated when evaluating UGPs (**Table S10, Figure 5D**). In general, we found that increasing the magnitude of the CV was concomitant with an increase in instances of IFF. This suggests that the use of FAIMS separation results in higher glycopeptide identifications that are false positives (i.e., those not originally in the sample), even when they provide unambiguous identifications via search algorithms. Evidence of IFF is an important limitation of FAIMS separation, given that it can (a) decrease the abundance of glycopeptides endogenously present in the sample, lowering their likelihood of detection, (b) result in confident identification of glycopeptides not present in the original sample, and (c) increase GSM and UGP counts through false-positive identifications. Thus, for accurate glycopeptide reporting, all FAIMS data should be evaluated thoroughly for these misidentifications.

## Conclusion

Glycoproteomic studies are usually accompanied by enrichment techniques to enhance detection and identification of glycosylated species, which can require many additional sample preparation steps. Further, each enrichment technique is often biased for particular glycan structures or peptide sequences. FAIMS separation offers the possibility to perform in-line separation based on ion mobility in the gas phase under an oscillating electric field. As FAIMS is a gas-phase separation technique, additional preparation steps are not required nor the losses associated with them, making it an appealing method to be used in glycoproteomic workflows. Here, we benchmarked FAIMS separation against traditional transmission techniques using three different samples: podocalyxin, a glycoprotein mixture, and a lectin enrichment of the human platelet sheddome.

In the case of a pure recombinant mucin-domain glycoprotein, we do not recommend exclusively performing FAIMS experiments, given the reduction in the identified glycosylated species and localized glycosites, along with the necessity for higher sample injections. However, in the case of more complex samples, FAIMS improved the glycoproteomic coverage detected, complementing results from traditional transmission techniques, while also lowering apparent polymer contamination. That said, FAIMS introduced a two- to five-fold increase in fragmentation between ionization and detection, which can reduce the detection of endogenous peptides while generating false identifications. However, even after removing instances of IFF, FAIMS separation of the platelet glyco-sheddome sample at -40V still resulted in a larger number of glycopeptide identifications. Taken together, FAIMS separation can improve identification of glycopeptides in complex samples but requires additional validation steps to ensure accurate reporting.

## Materials and Methods

### Materials

Unless otherwise mentioned, solutions were made using LCMS grade water (Pierce, 85178), acetonitrile (ACN, Honeywell, LC015), formic acid (Pierce, 85178), and ammonium bicarbonate (AmBic, Honeywell Fluka, 40867).

#### Sample preparation

##### Podocalyxin

Aliquots containing 10 µg of podocalyxin (R&D Systems, 1658-PD) recombinantly expressed in NS0 cells were reconstituted at a concentration of 1 µg/µL in 50 mM AmBic and subjected to different mucinase treatments. The first aliquot was digested using 0.2 µg of BT4244^28^ and cotreated with 3 U SialEXO (Genovis, G1-SM1-020) overnight at 37 °C; the second was digested using 0.1 µg of SmE^27^. The samples were then diluted to 20 µL using 50 mM AmBic. After digestion, the first aliquot was further treated with 10 U of PNGaseF (New England BioLabs, P0705S) for 8 hr at 37 °C.

##### Protein Mixture

The protein mixture was created by combining equal masses (5 µg) of each protein. Coagulation factor XII (Invitrogen, RP-43076), von Willebrand factor (Invitrogen, RP-43132), transferrin (R&D systems, 2914-HT), and apolipoprotein E (MoleCular Innovations, HaPo-E-5181) were isolated from human plasma. Bovine alpha-2-hs-glycoprotein was isolated from plasma (Fetuin, Promega, V4961), and monocyte differentiation antigen CD14 was recombinantly expressed in CHO cells (R&D systems, 383-CD-050/CF).

##### WGA enriched platelets

Expired apheresis platelets (BloodWorks Northwest, 2965-02) were enriched using wheat germ agglutinin (WGA) using previously described methods^31^. In short, seven apheresis units were washed and lysed using sonication. The supernatant was collected and enriched using a WGA agarose (Vector Laboratories, AL-1023) gravity flow column. After elution, the sample was concentrated using a centrifugal filter (Amicon, UCF910024), buffer exchanged into 50 mM AmBic pH 7.5, frozen, and shipped on dry ice. Fifty micrograms of the sample were digested by adding SmE to a ratio of 1:20 and incubating at 37 °C for 6 hours.

### Tryptic digestion

Dithiothreitol (DTT; Sigma, D0632) was added to a final concentration of 2 mM and the sample was reduced for 20 min at 65 °C. After allowing the sample to cool to room temperature, the sample was alkylated using iodoacetamide (IAA; Sigma, D0632) at a concentration of 3 mM and allowing it to react for 15 min in the dark at room temperature. After reduction and alkylation, sequencing-grade trypsin (Promega, V5117) was added to each sample at a ratio of 1:20 and allowed to digest overnight at 37 °C.

### Sample desalting

Strata X gravity flow columns (Phenomenex, 8B-S100-AAK) were activated using 1 mL of ACN followed by equilibration using 1 mL of 0.1% formic acid in water. Samples were acidified by addition of 1 µL of formic acid, diluted to a volume of 200 µL, and loaded onto the columns. After rinsing twice with 200 µL of 0.1% formic acid, the column was transferred to a second tube for elution with two additions of 150 µL of 40% ACN, 0.1% formic acid in water. Samples were dried via vacuum concentration and resuspended to a concentration of 300 ng/µL.

### MS Analysis

Samples were analyzed by online nanoflow liquid chromatography-tandem mass spectrometry using an Orbitrap Eclipse Tribrid mass spectrometer (Thermo Fisher Scientific) coupled to a Dionex UltiMate 3000 HPLC (Thermo Fisher Scientific). Samples were injected onto an Acclaim PepMap 100 column packed with 2 cm of 5 µm C18 material (Thermo Fisher, 164564) using 0.1% formic acid in water (solvent A). Peptides were then separated on a 15 cm PepMap RSLC EASY-Spray C18 column packed with 2 µm C18 material (Thermo Fisher, ES904) using a gradient from 0-35% solvent B (0.1% formic acid with 80% acetonitrile) in 60 min with a constant flow rate of 300 nL/min.

Full MS scans (MS1) were acquired in centroid mode in the Orbitrap at 120,000 resolution with a scan range of 300-1500 m/z. Ions were selected for fragmentation in order of decreasing intensity in a time-dependent manner, with a total cycle time of 3 seconds. Dynamic exclusion settings allowed 3 repeats in 10 seconds before exclusion for 10 seconds.

Fragmentation scans were collected in the Orbitrap at 15,000 resolution. For higher energy collisional dissociation (HCD) scans, ions were injected for a maximum of 50 ms, followed by fragmentation at 28% normalized collision energy (NCE). An EThcD scan was triggered if the HCD scan contained 3 out of 8 oxonium fingerprint ions (126.055, 138.055, 144.07, 168.065, 186.076, 204.086, 274.092, and 292.103) at greater than 5% relative intensity. EThcD used calibrated charge-dependent ETD times, maximum injection time was set to 200 ms, and supplemental activation was set to 15% NCE.

For FAIMS experiments, the Thermo Scientific FAIMS Pro (FMS02-10001) interface was employed in high resolution mode. In multiple CV experiments, the 3 second total cycle time was split equally between the CVs. For acquisitions without FAIMS separation, 300 ng were injected for each analysis, while 750 ng were injected when FAIMS was applied.

#### Data analysis and validation

Podocalyxin MS data files were searched using O-Pair hosted in the MetaMorpheus suite, and the glycoprotein mixture and platelet-extracted proteins were searched using Byonic. MS data and search results can be retrieved from the PRIDE repository (PXD041217). Finally, we reanalyzed data acquired by Alagesan *et al*.^26^ (PXD038673) with Byonic and investigated instances of in-FAIMS fragmentation. The term “Depleted Human Serum” will be used to refer to this data throughout the manuscript.

### O-Pair

Searches were performed using the MetaMorpheus suite version 0.0.320.^39^ The podocalyxin protein sequence was downloaded from Uniprot (O00592). Here, the *O-glycopeptide search* using the default *O-glycan* database was used, which contains 12 O-glycan compositions. Protease digestion was set to *nonspecific* with 24 missed cleavages to accommodate the variable N-termini generated with mucinase digestion. Deamidation on Asn and oxidation on Met were set as variable modifications, and carbamidomethyl on Cys was set as a constant fixed modification.

### Byonic

Searches were performed using Byonic v4.0.12.^40^ For the glycoprotein mixture, a database was created with the Uniprot protein sequences (P12763, P04275, P00748, P02787, P08571, P02649) and was used for all searches, while the human proteome (SwissProt, downloaded June 9, 2021) was used for the platelet-extracted and depleted human serum samples. All searches performed were fully tryptic, where the digestion termini were set to K/R on the C-terminus and maximum missed cleavages were set to 3. The fragmentation type was set to read directly from the scan headers. N- and O-glycopeptide searches were performed simultaneously, with the 9 most common O-glycans designated as common modifications with a maximum of two per peptide and 132 N-glycans designated as uncommon modifications with a maximum of one per peptide. The MS data of depleted human serum was searched using the same parameters, but only an O-glycan database. Other variable modifications (oxidation on Met and deamidation on Asn) were set as common for these searches, with a maximum of two common and one rare modification per peptide. Glycopeptides were filtered for score > 300 and log prob > 3 to ensure accurate peptide sequence.

### Manual Localization of O-glycopeptides

Manual validation of localized O-glycosites in podocalyxin was performed using Thermo XCalibur QualBrowser as previously described.^30^ In summary, glycan composition and peptide sequence were confirmed using the HCD spectrum. If the peptide had one possible O-glycosite in its sequence, O-glycans were localized to this site. If that was not the case, the EThcD spectrum was used to localize glycosites, where glycan masses were assigned based on the mass shifts on the c/z ions. If the presence of O-glycosylation could not be confirmed for the given site using the c/z fragment ions but its absence was confirmed for other probable sites, the site was designated as correct.^30^ Manual inspection was also performed for all peptides with length over 20 amino acids and with a charge state greater than 5.

### Calculations

A script to aid data analysis was written using RStudio (https://www.r-project.org/). Code for the calculation of aliphatic index was determined using the Peptides^41^ package, which assigned a value to each peptide depending on the number and type of aliphatic side chains in the peptide. Data visualization was performed with the ggplot2.^42^ Lastly, a two-sided student’s t-test in the rstatix^43^ package was used to determine whether there were significant differences in the average peptide length, aliphatic index, and glycan mass of peptides identified with and without FAIMS.

### Validation of in-source and in-FAIMS fragmentation

Glyco-SourceFragFinder was developed to identify potential cases of IFF/ISF, which were then manually validated. Glyco-SourceFragFinder has been made freely available at https://github.com/MalakerLab/Glyco-SourceFragFinder. Glyco-SourceFragFinder flagged identifications eluting within one minute of others with the same peptide backbone but bearing larger glycans

## Data availability

MS data acquired from reference ^26^ was deposited at PRIDE repository with the identifier PXD038673. MS files acquired and associated with this manuscript have been deposited at the PRIDE repository with the identifier (PXD041217), alongside output from search algorithms. PRIDE submission can currently be accessed with username “ “ and password “aRpK9yFj” Filtered and processed data is available as supplementary datasets. Glyco-SourceFragFinder is available as supplementary material and has been made freely available at https://github.com/MalakerLab/Glyco-SourceFragFinder.

## Author Contributions

S.A.M. conceptualization, supervision, project administration, funding acquisition; V.R.A., K.E.M., T.M.L, S.A.M. methodology; V.R.A., K.E.M. investigation; V.R.A. formal analysis, writing-original draft, visualization V.R.A, C.K. software, V.R.A, C.K., R.S validation; V.R.A., T.M.L., K.E.M., S.A.M., R.S. writing – review and editing.

## Acknowledgements

We thank Jeffrey Shabanowitz and Fanny Liu for their technical expertise and thoughtful conversations during the preparation of this manuscript. We also would like to acknowledge Carolyn Bertozzi and Judy Shon for the BT4244 and SmE plasmid, and Carolyn Bertozzi, Marie Hollenhorst, and Katherine Tiemeyer for providing the platelet glyco-sheddome sample. Finally, the authors would like to thank Alexandra Steigmeyer, Sarah Lowery, Deniz Ince, Vincent Chang, and Joann Chongsaritsinsuk for critical reading of this manuscript.

## References

(1) Wang, Y.-C.; Peterson, S. E.; Loring, J. F. Protein Post-Translational Modifications and Regulation of Pluripotency in Human Stem Cells. Cell Res 2014, 24 (2), 143–160. https://doi.org/10.1038/cr.2013.151.

(2) An, H. J.; Froehlich, J. W.; Lebrilla, C. B. Determination of Glycosylation Sites and Site-Specific Heterogeneity in Glycoproteins. Curr Opin Chem Biol 2009, 13 (4), 421–426. https://doi.org/10.1016/j.cbpa.2009.07.022.

(3) Stanley, P.; Moremen, K. W.; Lewis, N. E.; Taniguchi, N.; Aebi, M. N-Glycans. In Essentials of Glycobiology; Varki, A., Cummings, R. D., Esko, J. D., Stanley, P., Hart, G. W., Aebi, M., Mohnen, D., Kinoshita, T., Packer, N. H., Prestegard, J. H., Schnaar, R. L., Seeberger, P. H., Eds.; Cold Spring Harbor Laboratory Press: Cold Spring Harbor (NY), 2022.

(4) Rangel-Angarita, V.; Malaker, S. A. Mucinomics as the Next Frontier of Mass Spectrometry. ACS Chem. Biol. 2021, 16 (10), 1866–1883. https://doi.org/10.1021/acschembio.1c00384.

(5) Ince, D.; Lucas, T. M.; Malaker, S. A. Current Strategies for Characterization of Mucin-Domain Glycoproteins. Current Opinion in Chemical Biology 2022, 69, 102174. https://doi.org/10.1016/j.cbpa.2022.102174.

(6) Vigerust, D. J.; Shepherd, V. L. Virus Glycosylation: Role in Virulence and Immune Interactions. Trends in Microbiology 2007, 15 (5), 211–218. https://doi.org/10.1016/j.tim.2007.03.003.

(7) Sztain, T.; Ahn, S.-H.; Bogetti, A. T.; Casalino, L.; Goldsmith, J. A.; Seitz, E.; McCool, R. S.; Kearns, F. L.; Acosta-Reyes, F.; Maji, S.; Mashayekhi, G.; McCammon, J. A.; Ourmazd, A.; Frank, J.; McLellan, J. S.; Chong, L. T.; Amaro, R. E. A Glycan Gate Controls Opening of the SARS-CoV-2 Spike Protein. Nat. Chem. 2021, 13 (10), 963–968. https://doi.org/10.1038/s41557-021-00758-3.

(8) Lucas, T. M.; Gupta, C.; Altman, M. O.; Sanchez, E.; Naticchia, M. R.; Gagneux, P.; Singharoy, A.; Godula, K. Mucin-Mimetic Glycan Arrays Integrating Machine Learning for Analyzing Receptor Pattern Recognition by Influenza A Viruses. Chem 2021, 7 (12), 3393–3411. https://doi.org/10.1016/j.chempr.2021.09.015.

(9) Reily, C.; Stewart, T. J.; Renfrow, M. B.; Novak, J. Glycosylation in Health and Disease. Nat Rev Nephrol 2019, 15 (6), 346–366. https://doi.org/10.1038/s41581-019-0129-4.

(10) Dashti, H.; Pabon Porras, M. A.; Mora, S. Glycosylation and Cardiovascular Diseases. In The Role of Glycosylation in Health and Disease; Lauc, G., Trbojević-Akmačić, I., Eds.; Advances in Experimental Medicine and Biology; Springer International Publishing: Cham, 2021; pp 307–319. https://doi.org/10.1007/978-3-030-70115-4_15.

(11) Pereira, M. S.; Alves, I.; Vicente, M.; Campar, A.; Silva, M. C.; Padrão, N. A.; Pinto, V.; Fernandes, Â.; Dias, A. M.; Pinho, S. S. Glycans as Key Checkpoints of T Cell Activity and Function. Frontiers in Immunology 2018, 9, 2754. https://doi.org/10.3389/fimmu.2018.02754.

(12) Riley, N. M.; Bertozzi, C. R.; Pitteri, S. J. A Pragmatic Guide to Enrichment Strategies for Mass Spectrometry–Based Glycoproteomics. Mol Cell Proteomics 2020, 20, 100029. https://doi.org/10.1074/mcp.R120.002277.

(13) Li, H.; Bendiak, B.; Siems, W. F.; Gang, D. R.; Hill, H. H. Determining the Isomeric Heterogeneity of Neutral Oligosaccharide-Alditols of Bovine Submaxillary Mucin Using Negative Ion Traveling Wave Ion Mobility Mass Spectrometry. Anal. Chem. 2015, 87 (4), 2228–2235. https://doi.org/10.1021/ac503754k.

(14) Fenn, L. S.; McLean, J. A. Structural Separations by Ion Mobility-MS for Glycomics and Glycoproteomics. Methods Mol Biol 2013, 951, 171–194. https://doi.org/10.1007/978-1-62703-146-2_12.

(15) Fenn, L. S.; McLean, J. A. Simultaneous Glycoproteomics on the Basis of Structure Using Ion Mobility-Mass Spectrometry. Mol. BioSyst. 2009, 5 (11), 1298–1302. https://doi.org/10.1039/B909745G.

(16) Liu, F. C.; Cropley, T. C.; Ridgeway, M. E.; Park, M. A.; Bleiholder, C. Structural Analysis of the Glycoprotein Complex Avidin by Tandem-Trapped Ion Mobility Spectrometry–Mass Spectrometry (Tandem-TIMS/MS). Anal. Chem. 2020, 92 (6), 4459–4467. https://doi.org/10.1021/acs.analchem.9b05481.

(17) Pu, Y.; Ridgeway, M. E.; Glaskin, R. S.; Park, M. A.; Costello, C. E.; Lin, C. Separation and Identification of Isomeric Glycans by Selected Accumulation-Trapped Ion Mobility Spectrometry-Electron Activated Dissociation Tandem Mass Spectrometry. Anal. Chem. 2016, 88 (7), 3440–3443. https://doi.org/10.1021/acs.analchem.6b00041.

(18) Liu, F. C.; Kirk, S. R.; Caldwell, K. A.; Pedrete, T.; Meier, F.; Bleiholder, C. Tandem Trapped Ion Mobility Spectrometry/Mass Spectrometry (TTIMS/MS) Reveals Sequence-Specific Determinants of Top-Down Protein Fragment Ion Cross Sections. Anal. Chem. 2022, 94 (23), 8146–8155. https://doi.org/10.1021/acs.analchem.1c05171.

(19) Mookherjee, A.; Guttman, M. Bridging the Structural Gap of Glycoproteomics with Ion Mobility Spectrometry. Current Opinion in Chemical Biology 2018, 42, 86–92. https://doi.org/10.1016/j.cbpa.2017.11.012.

(20) Hebert, A. S.; Prasad, S.; Belford, M. W.; Bailey, D. J.; McAlister, G. C.; Abbatiello, S. E.; Huguet, R.; Wouters, E. R.; Dunyach, J.-J.; Brademan, D. R.; Westphall, M. S.; Coon, J. J. Comprehensive Single-Shot Proteomics with FAIMS on a Hybrid Orbitrap Mass Spectrometer. Anal. Chem. 2018, 90 (15), 9529–9537. https://doi.org/10.1021/acs.analchem.8b02233.

(21) Swearingen, K. E.; Moritz, R. L. High Field Asymmetric Waveform Ion Mobility Spectrometry (FAIMS) for Mass Spectrometry-Based Proteomics. Expert Rev Proteomics 2012, 9 (5), 505–517. https://doi.org/10.1586/epr.12.50.

(22) Pfammatter, S.; Bonneil, E.; McManus, F. P.; Prasad, S.; Bailey, D. J.; Belford, M.; Dunyach, J.-J.; Thibault, P. A Novel Differential Ion Mobility Device Expands the Depth of Proteome Coverage and the Sensitivity of Multiplex Proteomic Measurements *. Molecular &Cellular Proteomics 2018, 17 (10), 2051–2067. https://doi.org/10.1074/mcp.TIR118.000862.

(23) Ahmad Izaham, A. R.; Ang, C.-S.; Nie, S.; Bird, L. E.; Williamson, N. A.; Scott, N. E. What Are We Missing by Using Hydrophilic Enrichment? Improving Bacterial Glycoproteome Coverage Using Total Proteome and FAIMS Analyses. J. Proteome Res. 2021, 20 (1), 599–612. https://doi.org/10.1021/acs.jproteome.0c00565.

(24) Fang, P.; Ji, Y.; Silbern, I.; Viner, R.; Oellerich, T.; Pan, K.-T.; Urlaub, H. Evaluation and Optimization of High-Field Asymmetric Waveform Ion-Mobility Spectrometry for Multiplexed Quantitative Site-Specific N-Glycoproteomics. Anal. Chem. 2021, 93 (25), 8846–8855. https://doi.org/10.1021/acs.analchem.1c00802.

(25) Chandler, K. B.; Marrero Roche, D. E.; Sackstein, R. Multidimensional Separation and Analysis of Alpha-1-Acid Glycoprotein N-Glycopeptides Using High-Field Asymmetric Waveform Ion Mobility Spectrometry (FAIMS) and Nano-Liquid Chromatography Tandem Mass Spectrometry. Anal Bioanal Chem 2022. https://doi.org/10.1007/s00216-022-04435-3.

(26) Alagesan, K.; Ahmed-Begrich, R.; Charpentier, E. Improved N- and O-Glycopeptide Identification Using High-Field Asymmetric Waveform Ion Mobility Spectrometry (FAIMS); preprint; Biochemistry, 2022. https://doi.org/10.1101/2022.12.12.520086.

(27) Chongsaritsinsuk, J.; Steigmeyer, A. D.; Mahoney, K. E.; Rosenfeld, M. A.; Lucas, T. M.; Ince, D.; Kearns, F. L.; Battison, A. S.; Hollenhorst, M. A.; Shon, D. J.; Tiemeyer, K. H.; Attah, V.; Kwon, C.; Bertozzi, C. R.; Ferracane, M. J.; Amaro, R. E.; Malaker, S. A. Glycoproteomic Landscape and Structural Dynamics of TIM Family Immune Checkpoints Enabled by Mucinase SmE. bioRxiv February 3, 2023, p 2023.02.01.526488. https://doi.org/10.1101/2023.02.01.526488.

(28) Shon, D. J.; Malaker, S. A.; Pedram, K.; Yang, E.; Krishnan, V.; Dorigo, O.; Bertozzi, C. R. An Enzymatic Toolkit for Selective Proteolysis, Detection, and Visualization of Mucin-Domain Glycoproteins. Proc Natl Acad Sci U S A 2020, 117 (35), 21299–21307. https://doi.org/10.1073/pnas.2012196117.

(29) Syka, J. E. P.; Coon, J. J.; Schroeder, M. J.; Shabanowitz, J.; Hunt, D. F. Peptide and Protein Sequence Analysis by Electron Transfer Dissociation Mass Spectrometry. Proceedings of the National Academy of Sciences 2004, 101 (26), 9528–9533. https://doi.org/10.1073/pnas.0402700101.

(30) Rangel-Angarita, V.; Mahoney, K. E.; Ince, D.; Malaker, S. A. A Systematic Comparison of Current Bioinformatic Tools for Glycoproteomics Data. bioRxiv March 18, 2022, p x2022.03.15.484528. https://doi.org/10.1101/2022.03.15.484528.

(31) Hollenhorst, M. A.; Tiemeyer, K. H.; Mahoney, K. E.; Aoki, K.; Ishihara, M.; Lowery, S. C.; Rangel-Angarita, V.; Bertozzi, C. R.; Malaker, S. A. Comprehensive Analysis of Platelet Glycoprotein Ibα Ectodomain Glycosylation. Journal of Thrombosis and Haemostasis 2023. https://doi.org/10.1016/j.jtha.2023.01.009.

(32) Bagdonaite, I.; Malaker, S. A.; Polasky, D. A.; Riley, N. M.; Schjoldager, K.; Vakhrushev, S. Y.; Halim, A.; Aoki-Kinoshita, K. F.; Nesvizhskii, A. I.; Bertozzi, C. R.; Wandall, H. H.; Parker, B. L.; Thaysen-Andersen, M.; Scott, N. E. Glycoproteomics. Nat Rev Methods Primers 2022, 2 (1), 1–29. https://doi.org/10.1038/s43586-022-00128-4.

(33) Campbell, M. T.; Glish, G. L. Fragmentation in the Ion Transfer Optics after Differential Ion Mobility Spectrometry Produces Multiple Artifact Monomer Peaks. International Journal of Mass Spectrometry 2018, 425, 47–54. https://doi.org/10.1016/j.ijms.2018.01.007.

(34) Bleiholder, C.; Liu, F. C.; Chai, M. Comment on Effective Temperature and Structural Rearrangement in Trapped Ion Mobility Spectrometry. Anal. Chem. 2020, 92 (24), 16329–16333. https://doi.org/10.1021/acs.analchem.0c02052.

(35) Morsa, D.; Hanozin, E.; Eppe, G.; Quinton, L.; Gabelica, V.; Pauw, E. D. Effective Temperature and Structural Rearrangement in Trapped Ion Mobility Spectrometry. Anal. Chem. 2020, 92 (6), 4573–4582. https://doi.org/10.1021/acs.analchem.9b05850.

(36) Morsa, D.; Gabelica, V.; De Pauw, E. Fragmentation and Isomerization Due to Field Heating in Traveling Wave Ion Mobility Spectrometry. J. Am. Soc. Mass Spectrom. 2014, 25 (8), 1384–1393. https://doi.org/10.1007/s13361-014-0909-9.

(37) Liu, F. C.; Kirk, S. R.; Bleiholder, C. On the Structural Denaturation of Biological Analytes in Trapped Ion Mobility Spectrometry – Mass Spectrometry. Analyst 2016, 141 (12), 3722–3730. https://doi.org/10.1039/C5AN02399H.

(38) Merenbloom, S. I.; Flick, T. G.; Williams, E. R. How Hot Are Your Ions in TWAVE Ion Mobility Spectrometry? J Am Soc Mass Spectrom 2012, 23 (3), 553–562. https://doi.org/10.1007/s13361-011-0313-7.

(39) Lu, L.; Riley, N. M.; Shortreed, M. R.; Bertozzi, C. R.; Smith, L. M. O-Pair Search with MetaMorpheus for O-Glycopeptide Characterization. Nat Methods 2020, 17 (11), 1133–1138. https://doi.org/10.1038/s41592-020-00985-5.

(40) Bern, M.; Kil, Y. J.; Becker, C. Byonic: Advanced Peptide and Protein Identification Software. Curr Protoc Bioinformatics 2012, CHAPTER, Unit13.20. https://doi.org/10.1002/0471250953.bi1320s40.

(41) Osorio, D.; Rondon-Villareal, P.; Torres, R. Peptides: A Package for Data Mining of Antimicrobial Peptides. The R Journal 2015, 7 (1), 4–14.

(42) Wickham, H. Ggplot2: Elegant Graphics for Data Analysis, 978th-3rd-319th-24277th–4th ed.; Springer-Verlag: New York.

(43) Kassambara, A. Rstatix: Pipe-Friendly Framework for Basic Statistical Tests., 2023. https://rpkgs.datanovia.com/rstatix/.

