## Supplemental Information for "False positive glycopeptide identification via in-FAIMS fragmentation"

**This PDF file includes:**

Tables S1 to S9
Figures S1 to S13

**Additional supplementary materials not included in this document:**

Supplemental Dataset S1 – Podocalyxin SmE O-Pair output and manually validated peptides.
Supplemental Dataset S2 – Podocalyxin BTSia O-Pair output and calculated variables.
Supplemental Dataset S3 – Podocalyxin BTSia O-Pair manually validated peptides.
Supplemental Dataset S4 – Glycoprotein Mixture Byonic N-glycopeptide filtered output and calculated variables.
Supplemental Dataset S5 – Glycoprotein Mixture Byonic O-glycopeptide filtered output and calculated variables.
Supplemental Dataset S6 – Platelet sheddome sample Byonic N- and O- glycopeptide output.
Supplemental Dataset S7 – Plausible in-source fragmentation of Podocalyxin SmE and BTSia, platelet sheddome, and depleted human serum,[1] and manual validation.
Supplemental Code S1 – Matlab and R Studio scripts used to filter output results and calculate aliphatic index and average peptide length and glycan mass.

| Setting | GSMs | UGPs |
|---|---|---|

| | Counts | Percentage wrt No FAIMS | Counts | Percentage wrt No FAIMS |
|---|---|---|---|---|
| **No FAIMS** | 2611 | NA | 511 | NA |
| **-25CV** | 161 | 6% | 6 | 1% |
| **-30CV** | 602 | 23% | 43 | 8% |
| **-35CV** | 1070 | 41% | 106 | 21% |
| **-40CV** | 1256 | 48% | 159 | 31% |
| **-45CV** | 1271 | 49% | 244 | 48% |
| **-50CV** | 1290 | 49% | 269 | 53% |
| **-55CV** | 1073 | 41% | 255 | 50% |
| **-60CV** | 784 | 30% | 184 | 36% |
| **-65CV** | 491 | 19% | 101 | 20% |
| **-70CV** | 353 | 14% | 67 | 13% |
| **-75CV** | 178 | 7% | 34 | 7% |
| **-80CV** | 56 | 2% | 16 | 3% |

**Table S1. Percentage of identified UGPs and GSMs with respect to control experiments.** Recombinant podocalyxin was digested with mucinase SmE[2] and trypsin, followed by FAIMS separation and data analysis. RAW data was searched with O-pair hosted in the MetaMorpheus suite.[3] Glycopeptide spectral matches (GSMs) and unique glycopeptides (UGPs) were counted for each setting and are displayed in the "counts" section of the table. Finally, the fold change of each FAIMS setting was calculated with respect to (wrt) experiments without FAIMS, as shown in the "Percentage wrt No FAIMS" columns.

| Setting | GSMs | UGPs |
|---|---|---|

| | Counts | Percentage wrt No FAIMS | Counts | Percentage wrt No FAIMS |
|---|---|---|---|---|
| **No FAIMS** | 309 | NA | 110 | NA |
| **-25** | 1 | 0% | 1 | 1% |
| **-30** | 47 | 15% | 26 | 24% |
| **-35** | 95 | 31% | 36 | 33% |
| **-40** | 144 | 47% | 59 | 54% |
| **-45** | 152 | 49% | 61 | 55% |
| **-50** | 140 | 45% | 54 | 49% |
| **-55** | 113 | 37% | 40 | 36% |
| **-60** | 122 | 39% | 37 | 34% |
| **-65** | 76 | 25% | 21 | 19% |
| **-70** | 51 | 17% | 14 | 13% |
| **-75** | 15 | 5% | 5 | 5% |
| **-80** | 5 | 2% | 1 | 1% |
| **-85** | 0 | 0% | 0 | 0% |

**Table S2. Percentage of identified UGPs and GSMs with respect to control experiments.** Recombinant podocalyxin was digested with mucinase BT4244[4], sialidase and PNGaseF and trypsin, followed by FAIMS separation and data analysis. RAW data was searched with O-pair hosted in the MetaMorpheus suite.[3] Glycopeptide spectral matches (GSMs) and unique glycopeptides (UGPs) were counted for each setting and are displayed in the “counts” section of the table. Finally, the fold change of each FAIMS setting was calculated with respect to (wrt) experiments without FAIMS, as shown in the “Fold Change” columns.

| N-glycopeptides | | | | |
|---|---|---|---|---|
| **Setting** | **GSMs** | | **UGPs** | |
| | **Counts** | **Percentage wrt No FAIMS** | **Counts** | **Percentage wrt No FAIMS** |
| **No FAIMS** | 154 | NA | 39 | NA |
| **-20** | 0 | 0% | 0 | 0% |
| **-25** | 15 | 10% | 5 | 13% |
| **-30** | 28 | 18% | 7 | 18% |
| **-35** | 97 | 63% | 25 | 64% |
| **-40** | 135 | 88% | 28 | 72% |
| **-45** | 164 | 106% | 34 | 87% |
| **-50** | 166 | 108% | 32 | 82% |
| **-55** | 123 | 80% | 24 | 62% |
| **-60** | 35 | 23% | 13 | 33% |
| **-65** | 21 | 14% | 9 | 23% |
| **-70** | 19 | 12% | 7 | 18% |
| **-75** | 20 | 13% | 6 | 15% |
| **-80** | 7 | 5% | 4 | 10% |
| **-85** | 0 | 0% | 0 | 0% |
| **-40/-45** | 227 | 147% | 44 | 113% |
| **-40/-50** | 193 | 125% | 38 | 97% |
| **-25/-30/-35** | 80 | 52% | 20 | 51% |
| **-40/-45/-50** | 295 | 192% | 37 | 95% |
| **-55/-65/-75** | 48 | 31% | 18 | 46% |

**Table S3. Percentage of identified N- UGPs and GSMs with respect to control experiments.** A mixture of six glycoproteins was generated, digested using trypsin, and subjected to FAIMS followed by MS analysis. Resulting RAW data was searched with Byonic. N-GSMs and UGPs were counted for each setting and are displayed in the "counts" section of the table. Finally, the fold change of each FAIMS setting was calculated with respect to the no FAIMS experiment, shown in the "Percentage wrt No FAIMS" columns.

| O-glycopeptides | | | | |
|---|---|---|---|---|
| **Setting** | **GSMs** | | **UGPs** | |
| | **Counts** | **Percentage wrt No FAIMS** | **Counts** | **Percentage wrt No FAIMS** |
| **No FAIMS** | 189 | NA | 25 | NA |
| **-20** | 5 | 3% | 1 | 4% |
| **-25** | 28 | 15% | 6 | 24% |
| **-30** | 37 | 20% | 7 | 28% |
| **-35** | 117 | 62% | 15 | 60% |
| **-40** | 171 | 90% | 24 | 96% |
| **-45** | 203 | 107% | 28 | 112% |
| **-50** | 141 | 75% | 24 | 96% |
| **-55** | 90 | 48% | 13 | 52% |
| **-60** | 32 | 17% | 6 | 24% |
| **-65** | 11 | 6% | 3 | 12% |
| **-70** | 0 | 0% | 0 | 0% |
| **-75** | 0 | 0% | 0 | 0% |
| **-80** | 0 | 0% | 0 | 0% |
| **-85** | 0 | 0% | 0 | 0% |
| **-40/-45** | 284 | 150% | 23 | 92% |
| **-40/-50** | 206 | 109% | 25 | 100% |
| **-25/-30/-35** | 122 | 65% | 13 | 52% |
| **-40/-45/-50** | 252 | 133% | 25 | 100% |
| **-55/-65/-75** | 44 | 23% | 11 | 44% |

**Table S4. Percentage of identified O- UGPs and GSMs with respect to control experiments.** A mixture of six glycoproteins was generated, digested using trypsin, and subjected to FAIMS followed by MS analysis. Resulting RAW data was searched with Byonic. N-GSMs and UGPs were counted for each setting and are displayed in the “counts” section of the table. Finally, the fold change of each FAIMS setting was calculated with respect to the no FAIMS experiment, shown in the “Percentage wrt No FAIMS” columns.

| Type | Setting | Code output | | Flagged but not ISF/IFF | Confirmed ISF/IFF | Confirmed percentage of ISF/IFF (%) |
|---|---|---|---|---|---|---|
| | | Not ISF/IFF | Flagged as ISF/IFF | | | |
| GSMs | -40V | 885 | 350 | 160 | 190 | 15.4 |
| | -45V | 796 | 426 | 132 | 294 | 24.1 |
| | -50V | 753 | 460 | 177 | 283 | 23.3 |
| | No FAIMS | 1827 | 738 | 550 | 188 | 7.3 |
| UGPs | -40V | 299 | 107 | 59 | 48 | 11.8 |
| | -45V | 280 | 129 | 50 | 79 | 19.3 |
| | -50V | 249 | 140 | 62 | 78 | 20.1 |
| | No FAIMS | 528 | 203 | 159 | 44 | 6.0 |

**Table S5. ISF and IFF observed in SmE-digested podocalyxin.** Recombinant podocalyxin was digested with mucinase SmE[2] and trypsin, followed by FAIMS separation and data analysis. RAW data was searched with O-pair, hosted in the MetaMorpheus suite.[3] GSMs and UGPs were counted for each setting and are displayed in in the table. Glyco-SourceFragFinder was used to flag identifications that eluted within a minute of another identification from the same glycopeptide backbone sequence with a larger glycan structure attached to it, delimited as “Flagged as ISF/IFF”. Those that were not flagged are delimited as “Not ISF/IFF”. Following, flagged ISF/IFF identifications were validated by extracting the chromatographic profiles, as well as those from identifications with the same glycopeptide backbone sequence with a larger glycan structure eluting within a minute of it. If the two chromatographic profiles aligned, it was confirmed as ISF/IFF. Finally, the percentage of ISF/IFF was calculated with respect to the total number of GSMs and UGPs.

| Type | Setting | Code output | | Flagged but not ISF/IFF | Confirmed ISF/IFF | Confirmed percentage of ISF/IFF (%) |
|---|---|---|---|---|---|---|
| | | Not ISF/IFF | Flagged as ISF/IFF | | | |
| GSMs | -40V | 112 | 32 | 19 | 13 | 9.0 |
| | -45V | 108 | 44 | 12 | 32 | 21.1 |
| | -50V | 89 | 51 | 13 | 38 | 27.1 |
| | No FAIMS | 237 | 72 | 59 | 13 | 4.2 |
| UGPs | -40V | 49 | 15 | 8 | 7 | 10.9 |
| | -45V | 47 | 15 | 4 | 11 | 17.7 |
| | -50V | 39 | 19 | 6 | 13 | 22.4 |
| | No FAIMS | 84 | 31 | 23 | 8 | 7.0 |

**Table S6. IFF and ISF observed in BT4244-digested podocalyxin.** Recombinant podocalyxin was digested with mucinase BT4244[4], sialidase and PNGaseF and trypsin, followed by FAIMS separation and data analysis. RAW data was searched with O-pair, hosted in the MetaMorpheus suite.[3] GSMs and UGPs were counted for each setting and are displayed in in the table. Glyco-SourceFragFinder was used to flag identifications that eluted within a minute of another identification from the same glycopeptide backbone sequence with a larger glycan structure attached to it, delimited as “Flagged as ISF/IFF”. Those that were not flagged are delimited as “Not ISF/IFF”. Following, flagged ISF/IFF identifications were validated by extracting the chromatographic profiles, as well as those from identifications with the same glycopeptide backbone sequence with a larger glycan structure eluting within a minute of it. If the two chromatographic profiles aligned, it was confirmed as ISF/IFF. Finally, the percentage of ISF/IFF was calculated with respect to the total number of GSMs and UGPs.

| Type | Setting | Code output | | Flagged but not ISF/IFF | Confirmed ISF/IFF | Confirmed percentage of ISF (%) |
|---|---|---|---|---|---|---|
| | | Not ISF | Flagged as ISF | | | |
| GSMs | -40V | 254 | 195 | 141 | 54 | 12.0 |
| | -45V | 159 | 157 | 89 | 68 | 21.5 |
| | No FAIMS | 136 | 60 | 52 | 8 | 4.1 |
| UGPs | -40V | 90 | 46 | 30 | 16 | 11.8 |
| | -45V | 51 | 29 | 13 | 16 | 20.0 |
| | No FAIMS | 67 | 22 | 17 | 5 | 5.6 |

**Table S7. IFF and ISF observed the O-glycopeptide platelet sheddome.** The platelet sheddome was enriched using LAC, digested with mucinase SmE[2] and trypsin, followed by FAIMS separation and data analysis. RAW data was searched with O-pair, hosted in the MetaMorpheus suite.[3] GSMs and UGPs were counted for each setting and are displayed in in the table. Glyco-SourceFragFinder was used to flag identifications that eluted within a minute of another identification from the same glycopeptide backbone sequence with a larger glycan structure attached to it, delimited as “Flagged as ISF/IFF”. Those that were not flagged are delimited as “Not ISF/IFF”. Following, flagged ISF/IFF identifications were validated by extracting the chromatographic profiles, as well as those from identifications with the same glycopeptide backbone sequence with a larger glycan structure eluting within a minute of it. If the two chromatographic profiles aligned, it was confirmed as ISF/IFF. Finally, the percentage of ISF/IFF was calculated with respect to the total number of GSMs and UGPs.

| Type of identification | FAIMS Setting | | Code output | | Flagged but not ISF | Confirmed ISF | Confirmed percentage of ISF (%) |
|---|---|---|---|---|---|---|---|
| | CV | Replicate | Not ISF | Flagged as ISF | | | |
| GSMs | **-40V** | 1 | 141 | 100 | 33 | 65 | 27.0 |
| | **-40V** | 2 | 131 | 92 | 25 | 67 | 30.0 |
| | **-45V** | 1 | 147 | 112 | 29 | 83 | 32.0 |
| | **-45V** | 2 | 169 | 108 | 26 | 82 | 29.6 |
| | **-50V** | 1 | 140 | 94 | 18 | 76 | 32.5 |
| | **-50V** | 2 | 137 | 90 | 18 | 72 | 31.7 |
| UGPs | **-40V** | 1 | 101 | 66 | 23 | 43 | 25.4 |
| | **-40V** | 2 | 100 | 59 | 17 | 42 | 26.4 |
| | **-45V** | 1 | 111 | 77 | 17 | 60 | 31.9 |
| | **-45V** | 2 | 115 | 75 | 16 | 59 | 31.1 |
| | **-50V** | 1 | 100 | 62 | 14 | 48 | 29.6 |
| | **-50V** | 2 | 99 | 65 | 12 | 53 | 32.3 |

**Table S8. IFF observed in depleted human serum data acquired by Alagesan et. al.**[1] Depleted human serum RAW data, acquired in reference 1, was searched with Byonic[5] using the same settings applied for the platelet glyco-sheddome. Due to data availability, only settings -40V, -45V, and -50V were studied. These settings were taken in technical replicates. GSMs and UGPs were counted for each setting and are displayed in in the table. Glyco-SourceFragFinder was used to flag identifications that eluted within a minute of another identification from the same glycopeptide backbone sequence with a larger glycan structure attached to it, delimited as “Flagged as ISF/IFF”. Those that were not flagged are delimited as “Not ISF/IFF”. Following, flagged ISF/IFF identifications were validated, by extracting the chromatographic profiles, as well as those from identifications with the same glycopeptide backbone sequence with a larger glycan structure eluting within a minute of it. If the two chromatographic profiles aligned, it was confirmed as ISF/IFF. Finally, the percentage of ISF/IFF was calculated with respect to the total number of GSMs and UGPs.

<table>
<tr><th rowspan="2">Type of identification</th><th rowspan="2">Sample</th><th colspan="4">Percentage of total identifications from ISF/IFF (%)</th></tr>
<tr><th>-40V</th><th>-45V</th><th>-50V</th><th>No FAIMS</th></tr>
<tr><td rowspan="7">GSMs</td><td>Podocalyxin SmE</td><td>15.4</td><td>24.1</td><td>23.3</td><td>7.3</td></tr>
<tr><td>Podocalyxin BTSia</td><td>9</td><td>21.1</td><td>27.1</td><td>4.2</td></tr>
<tr><td>Platelet Sheddome</td><td>12</td><td>21.5</td><td>-</td><td>4.1</td></tr>
<tr><td>Depleted Human Serum Rep 1</td><td>27</td><td>32</td><td>32.5</td><td>-</td></tr>
<tr><td>Depleted Human Serum Rep 2</td><td>30</td><td>29.6</td><td>23.7</td><td>-</td></tr>
<tr><td>Average</td><td>18.7</td><td>25.7</td><td>28.7</td><td>5.2</td></tr>
<tr><td>Standard Deviation</td><td>9.3</td><td>4.9</td><td>4.3</td><td>1.8</td></tr>
<tr><td rowspan="7">UGPs</td><td>Podocalyxin SmE</td><td>11.8</td><td>19.3</td><td>20.1</td><td>6</td></tr>
<tr><td>Podocalyxin BTSia</td><td>10.9</td><td>17.7</td><td>22.4</td><td>7</td></tr>
<tr><td>Platelet Sheddome</td><td>11.8</td><td>20</td><td>-</td><td>5.6</td></tr>
<tr><td>Depleted Human Serum Rep 1</td><td>25.4</td><td>31.9</td><td>29.6</td><td>-</td></tr>
<tr><td>Depleted Human Serum Rep 2</td><td>26.4</td><td>31.1</td><td>32.3</td><td>-</td></tr>
<tr><td>Average</td><td>17.3</td><td>24</td><td>26.1</td><td>6.2</td></tr>
<tr><td>Standard Deviation</td><td>7.9</td><td>6.9</td><td>5.8</td><td>0.7</td></tr>
</table>

**Table S9. Average number of Identifications from ISF/IFF.** Glyco-SourceFragFinder was used to flag possible instances of ISF/IFF. Following, flagged ISF/IFF identifications were validated by extracting the chromatographic profile of the flagged identification, as well as those from glycopeptides with the same backbone sequence and a larger glycan structure eluting within a minute of it. If the two chromatographic profiles aligned, it was confirmed as ISF/IFF. Here, the percentage of identifications confirmed as ISF/IFF are depicted for the three samples analyzed, and the four experimental settings studied. Following, the average of these percentages was calculated, as well as the standard deviation. Since the FAIMS setting -50V was not collected for the platelet sheddome sample, a percentage is not depicted there. Similarly, data acquired without FAIMS was not available in reference 1 and therefore, it was not used here.

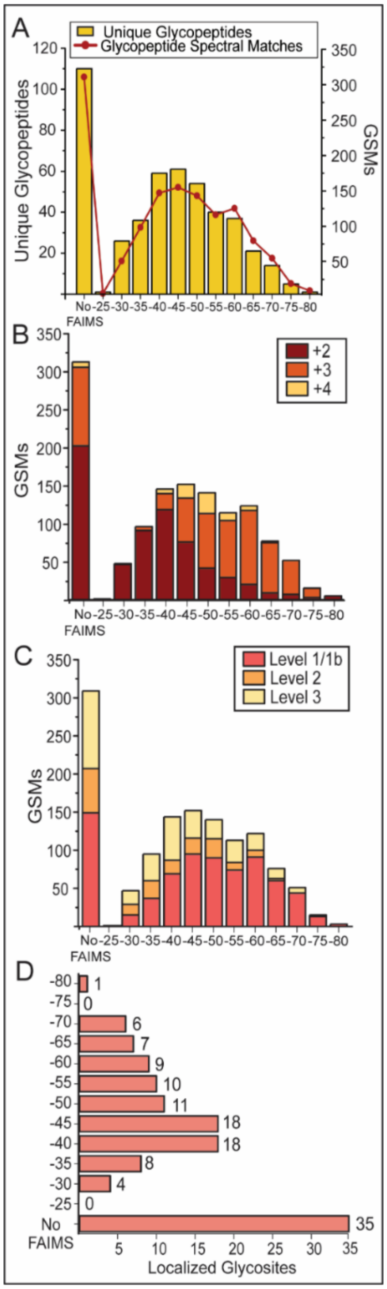


**Figure S1. Benchmarking FAIMS separation using BT4244 digested podocalyxin.** Recombinant podocalyxin was digested with mucinase BT4244[4], sialidase, PNGaseF, and trypsin, then subjected to FAIMS separation and MS analysis. Resulting MS data was searched with O-Pair. All figures were generated using OriginPro 2023. (A) GSMs (yellow, bar graph, right axis) and UGPs (red, line plot, left axis) identified with and without FAIMS. UGP lists were generated by removing duplicates of identical peptide sequences and glycan masses from search results. (B) Charge state distribution of different FAIMS settings. GSMs were classified by precursor charge state, and results were plotted as a stacked bar graph. Identified charge states range from +2 (dark red) to +5 (yellow). (C) GSMs were also classified based on the localization level assigned by O-Pair, where Level 1/1b (pink) represented fully localized GSMs, while Level 2 (orange) represented GSMs where at least one site could not be localized. Finally, Level 3 (yellow) illustrated GSMs where no glycan could be localized. (D) Following manual validation of GSMs, the number of localized glycosites per FAIMS setting was plotted.

| | Sample Size | Peptide Length | | Aliphatic Index | | Glycan Mass | |
|---|---|---|---|---|---|---|---|
| | | Mean | p-value | Mean | p-value | Mean | p-value |
| No FAIMS | 2565 | 12.5 | N/A | 52.1 | N/A | 1158.6 | N/A |
| -25 | 161 | 12.5 | | 47.6 | * | 890.0 | ** |
| -30 | 594 | 11.6 | ** | 54.6 | | 820.6 | ** |
| -35 | 1051 | 11.6 | ** | 59.1 | ** | 819.9 | ** |
| -40 | 1256 | 12.6 | | 56.7 | ** | 1016.4 | ** |
| -45 | 1271 | 13.3 | ** | 48.5 | ** | 1104.0 | * |
| -50 | 1290 | 13.9 | ** | 47.3 | ** | 1027.6 | ** |
| -55 | 1073 | 13.4 | ** | 46.4 | ** | 904.9 | ** |
| -60 | 784 | 13.2 | ** | 47.3 | ** | 778.4 | ** |
| -65 | 491 | 12.9 | ** | 46.3 | ** | 616.3 | ** |
| -70 | 353 | 13.4 | ** | 44.2 | ** | 584.5 | ** |
| -75 | 178 | 13.4 | ** | 42.8 | ** | 546.5 | ** |
| -80+ | 56 | 13.9 | N/A | 37.6 | N/A | 510.4 | N/A |

+ Sample size is too small to make a conclusion

* p < 0.05 — Value > No FAIMS
** p < 0.01 — Value < No FAIMS

**Figure S2. Properties of peptides preferentially separated in FAIMS in the podocalyxin SmE digest.** Average peptide length and aliphatic index were calculated using a program developed in the Scott lab.[6] A standardized two-sided student's t-test was performed against results from the experiment without FAIMS. Depicted are the sample size, average peptide length, aliphatic index, and total glycan mass per peptide for all CVs and the experiment without FAIMS. In the p-value column, empty cells denoted a lack of significance, whereas significance was indicated as * for p values <0.05 and ** for p values <0.01. N/A represented instances where the sample size was too small to perform a t-test. If the mean value of the tested variables (peptide length, aliphatic index, and glycan mass) at a given CV was found to be significantly higher when compared to experiments without FAIMS, it was denoted as red. Conversely, if it was found to be significantly lower, it was denoted as blue.

| | Sample Size | Peptide Length | | Aliphatic Index | | Glycan Mass | |
|---|---|---|---|---|---|---|---|
| | | Mean | p-value | Mean | p-value | Mean | p-value |
| No FAIMS | 309 | 11.3 | N/A | 61.6 | N/A | 620.9 | N/A |
| -25$^{+}$ | 1 | 22.0 | N/A | 9.1 | N/A | 365.1 | N/A |
| -30 | 47 | 12.9 | * | 64.9 | | 635.7 | |
| -35 | 95 | 11.8 | | 67.3 | | 553.1 | * |
| -40 | 144 | 11.1 | | 64.9 | | 544.5 | ** |
| -45 | 152 | 12.4 | * | 60.2 | | 574.4 | |
| -50 | 140 | 13.4 | ** | 55.3 | * | 600.3 | |
| -55 | 113 | 13.7 | ** | 54.8 | * | 540.6 | * |
| -60 | 122 | 13.7 | ** | 49.7 | ** | 556.4 | * |
| -65 | 76 | 14.3 | ** | 46.5 | ** | 509.4 | ** |
| -70 | 51 | 13.6 | ** | 48.9 | ** | 491.0 | ** |
| -75$^{+}$ | 15 | 13.3 | N/A | 47.2 | N/A | 510.5 | N/A |
| -80$^{+}$ | 5 | 7.0 | N/A | 84.3 | N/A | 203.1 | N/A |

$^{+}$ Sample size is too small to make a conclusion

* $p < 0.05$ — Value > No FAIMS (red)
** $p < 0.01$ — Value < No FAIMS (blue)

**Figure S3. Properties of peptides preferentially separated in FAIMS in the podocalyxin BT4244 digest.** Average peptide length and aliphatic index were calculated using a program developed in the Scott lab.[6] A standardized two-sided student's t-test was performed against results from the experiment without FAIMS. Depicted are the sample size, average peptide length, aliphatic index, and total glycan mass per peptide for all CVs and the experiment without FAIMS. In the p-value column, empty cells denoted a lack of significance, whereas significance was indicated as * for p values <0.05 and ** for p values <0.01. N/A represented instances where the sample size was too small to perform a t-test. If the mean value of the tested variables (peptide length, aliphatic index, and glycan mass) at a given CV was found to be significantly higher when compared to experiments without FAIMS, it was denoted as red. Conversely, if it was found to be significantly lower, it was denoted as blue.

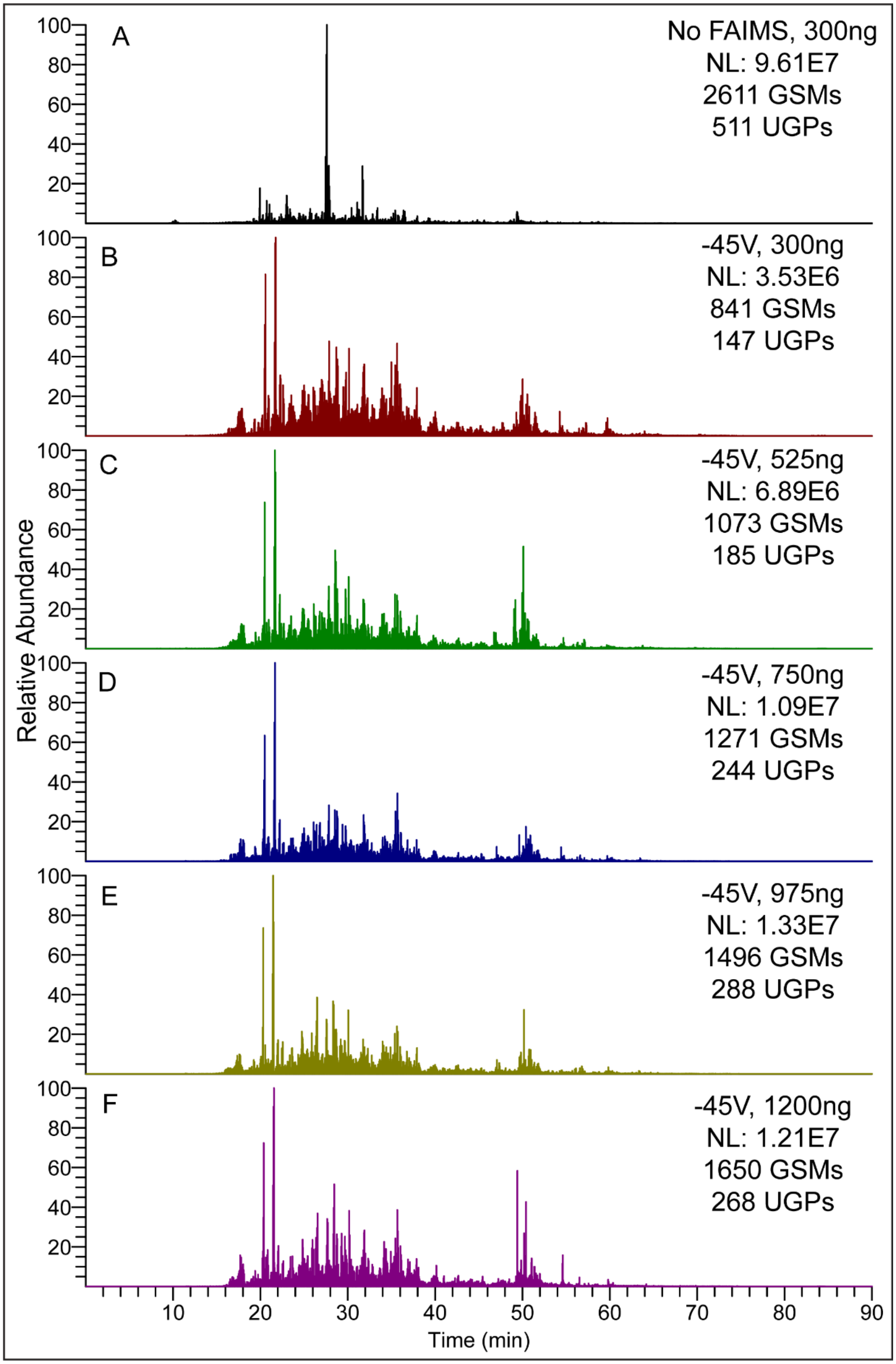


**Figure S4. Oxonium ion signal with increasing amounts of sample in -45V FAIMS separation.** Recombinant podocalyxin was digested with mucinase $SmE^2$ and trypsin. We performed a control experiment without FAIMS separation (A) and used CV -45V in FAIMS experiments using different sample amounts: 300 ng (B), 525 ng (C), 750 ng (D), 975 ng (E) and 1200 ng (F). The total ion chromatogram of m/z 204.0867 for each setting was shown, alongside the normalized abundance of each chromatogram.

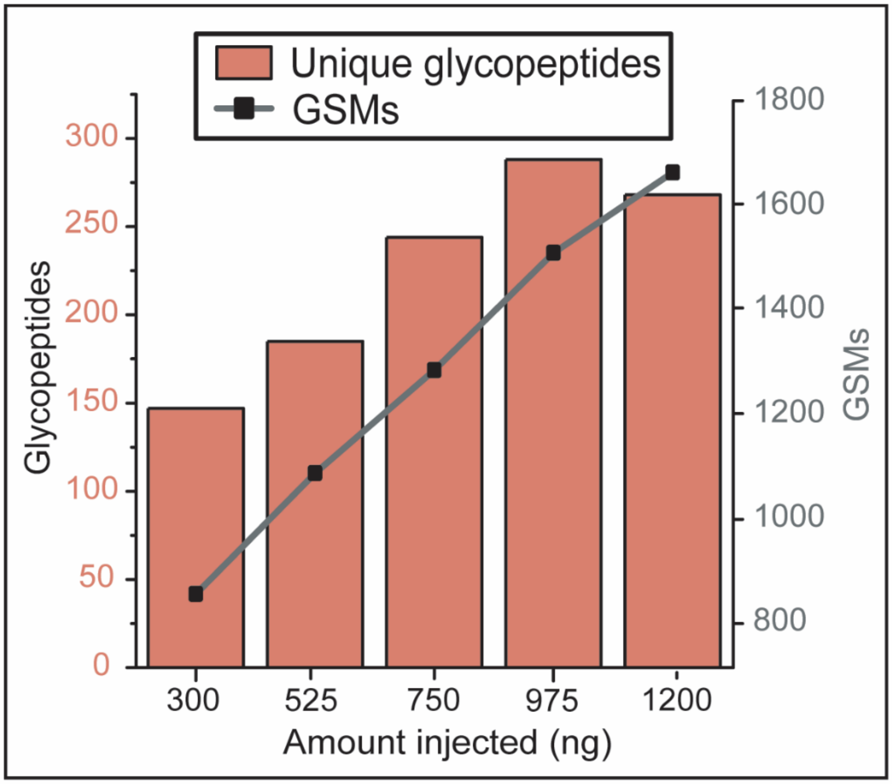


**Figure S5. Identified glycosylated species increase with injected sample amount.** Recombinant podocalyxin was digested with mucinase SmE$^2$ and trypsin, then subjected to FAIMS separation. Resulting RAW data was searched with O-Pair and results were analyzed. GSMs (gray, scatter plot) and UGPs (pink, bar graph) were analyzed for different sample amounts: 300 ng, 525 ng, 750 ng, 975 ng, and 1200 ng.

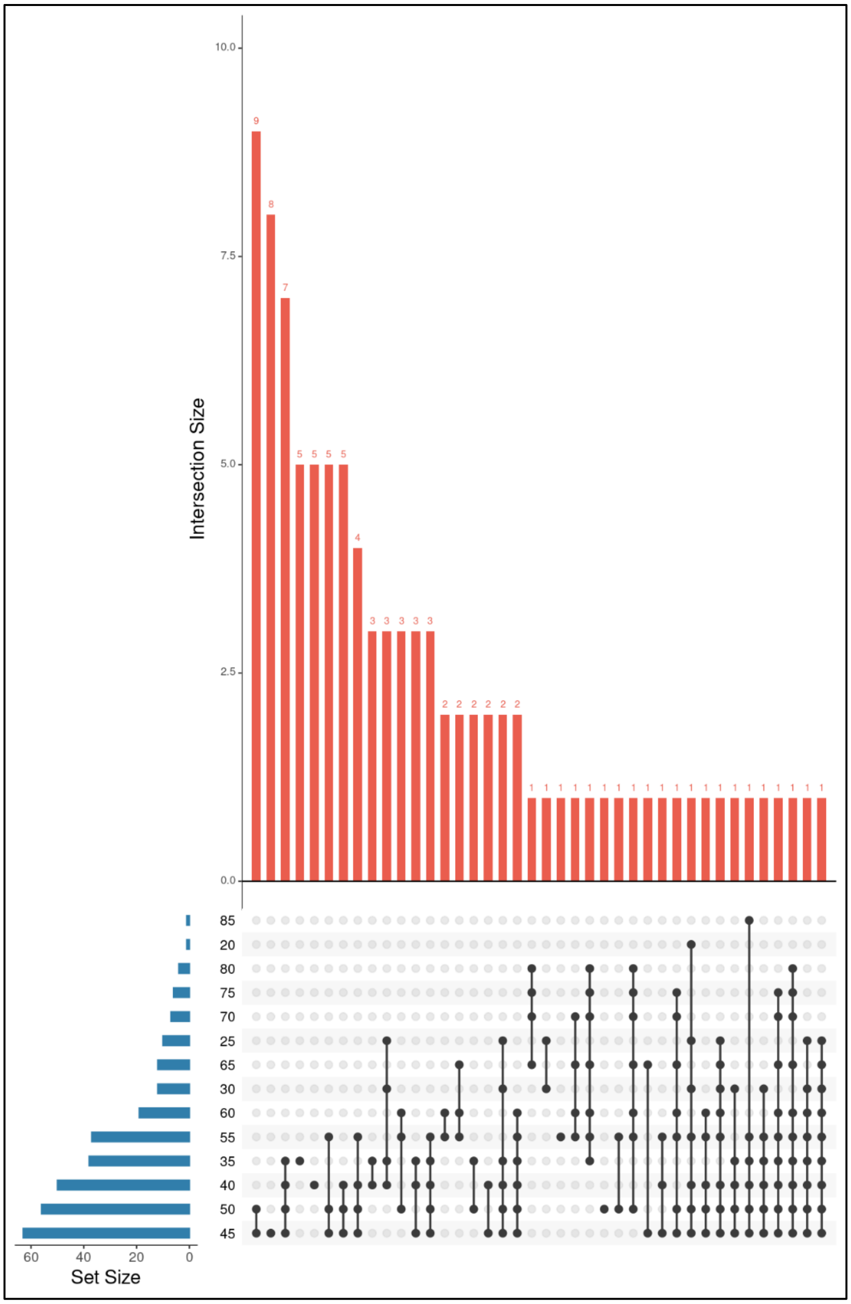


**Figure S6. Upset plot representing identified glycopeptide overlap among different CVs.** A mixture of six glycoproteins was generated, digested using trypsin, and subjected to FAIMS separation followed by MS analysis. The FAIMS CVs used are depicted on the left side of the graph, where the set size represented the number of UGPs identified in each. The overlap between FAIMS conditions is represented by the darkened dots and line connecting them, akin to the intersection that would be observed in a Venn diagram. The intersection size is depicted in the bar graphs. This figure was generated using Intervene.[7]

| | Sample Size | Peptide Length Mean | Peptide Length p-value | Aliphatic Index Mean | Aliphatic Index p-value | Glycan Mass Mean | Glycan Mass p-value |
|---|---|---|---|---|---|---|---|
| No FAIMS | 154 | 18.9 | N/A | 72.2 | N/A | 1981.8 | N/A |
| -25 $^{+}$ | 15 | 19.6 | N/A | 98.3 | N/A | 1880.1 | N/A |
| -30 | 28 | 20.3 | | 90.6 | ** | 1845.9 | |
| -35 | 97 | 20.6 | * | 85.0 | ** | 1920.7 | |
| -40 | 135 | 20.1 | | 72.4 | | 2079.9 | ** |
| -45 | 164 | 20.0 | | 63.2 | ** | 2101.2 | ** |
| -50 | 166 | 18.4 | | 68.8 | | 2038.6 | |
| -55 | 123 | 17.0 | ** | 62.9 | ** | 1982.3 | |
| -60 | 35 | 14.4 | ** | 60.9 | * | 1972.0 | |
| -65 $^{+}$ | 21 | 12.1 | N/A | 60.6 | N/A | 1962.2 | N/A |
| -70 $^{+}$ | 19 | 12.1 | N/A | 58.8 | N/A | 1917.4 | N/A |
| -75 $^{+}$ | 20 | 13.1 | N/A | 57.8 | N/A | 1917.5 | N/A |
| -80 $^{+}$ | 7 | 12.9 | N/A | 57.7 | N/A | 1903.4 | N/A |
| -40/-45 | 227 | 20.4 | * | 70.1 | | 2078.8 | * |
| -40/-50 | 193 | 20.3 | * | 67.8 | | 2089.2 | ** |
| -25/-30/-35 | 80 | 18.7 | | 85.3 | ** | 1922.0 | |
| -40/-45/-50 | 295 | 20.2 | * | 66.6 | | 2100.1 | ** |
| -55/-65/-75 | 48 | 16.0 | ** | 60.5 | * | 1786.0 | ** |

$^{+}$ Sample size is too small to make a conclusion

* $p < 0.05$   Value > No FAIMS

** $p < 0.01$   Value < No FAIMS

**Figure S7. Evaluation of identified N-glycopeptides features using FAIMS.** A mixture of six glycoproteins was generated, digested using trypsin, and subjected to FAIMS followed by MS analysis. Average peptide length and aliphatic index were calculated using a program developed in the Scott lab.[6] A standardized two-sided student's t-test was performed against results from the experiment without FAIMS. Depicted are the sample size, average peptide length, aliphatic index, and total glycan mass per peptide for all CVs and the experiment without FAIMS. In the p-value column, empty cells denoted a lack of significance, whereas significance was indicated as * for p values <0.05 and ** for p values <0.01. N/A represented instances where the sample size was too small to perform a t-test. If the mean value of the tested variables (peptide length, aliphatic index, and glycan mass) at a given CV was found to be significantly higher when compared to experiments without FAIMS, it was denoted as red. Conversely, if it was found to be significantly lower, it was denoted as blue.

| | Sample Size | Peptide Length | | Aliphatic Index | | Glycan Mass | |
|---|---|---|---|---|---|---|---|
| | | Mean | p-value | Mean | p-value | Mean | p-value |
| No FAIMS | 189 | 21.5 | N/A | 52.4 | N/A | 1160.8 | N/A |
| 20+ | 5 | 14.0 | N/A | 42.1 | N/A | 1312.0 | N/A |
| 25+ | 28 | 14.3 | N/A | 42.7 | N/A | 897.4 | N/A |
| 30 | 37 | 17.8 | ** | 47.1 | * | 917.2 | * |
| 35 | 117 | 22.2 | | 57.3 | ** | 1079.8 | |
| 40 | 171 | 23.9 | ** | 55.1 | * | 1262.1 | |
| 45 | 203 | 22.7 | | 52.5 | | 1313.7 | * |
| 50 | 141 | 20.1 | | 47.0 | ** | 1151.5 | |
| 55 | 90 | 16.4 | ** | 44.3 | ** | 858.0 | ** |
| 60 | 32 | 14.6 | ** | 42.9 | ** | 623.2 | ** |
| 65+ | 11 | 15.2 | N/A | 50.0 | N/A | 656.0 | N/A |
| 40/45 | 284 | 22.0 | | 55.7 | ** | 1142.1 | |
| 40/50 | 206 | 22.4 | | 53.0 | | 1233.2 | |
| 25/30/35 | 122 | 18.1 | ** | 51.2 | | 876.5 | ** |
| 40/45/50 | 252 | 22.3 | | 52.9 | | 1244.4 | |
| 55/65/75 | 44 | 14.9 | ** | 43.3 | ** | 836.3 | ** |

+Sample size is too small to make a conclusion

| | |
|---|---|
| * $p < 0.05$ | Value > No FAIMS (red) |
| ** $p < 0.01$ | Value < No FAIMS (blue) |

**Figure S8. Evaluation of identified O-glycopeptides features using FAIMS.** A mixture of six glycoproteins was generated, digested using trypsin, and subjected to FAIMS followed by MS analysis. Average peptide length and aliphatic index were calculated using a program developed in the Scott lab.[6] A standardized two-sided student's t-test was performed against results from the experiment without FAIMS. Depicted are the sample size, average peptide length, aliphatic index, and total glycan mass per peptide for all CVs and the experiment without FAIMS. In the p-value column, empty cells denoted a lack of significance, whereas significance was indicated as * for p values <0.05 and ** for p values <0.01. N/A represented instances where the sample size was too small to perform a t-test. If the mean value of the tested variables (peptide length, aliphatic index, and glycan mass) at a given CV was found to be significantly higher when compared to experiments without FAIMS, it was denoted as red. Conversely, if it was found to be significantly lower, it was denoted as blue.

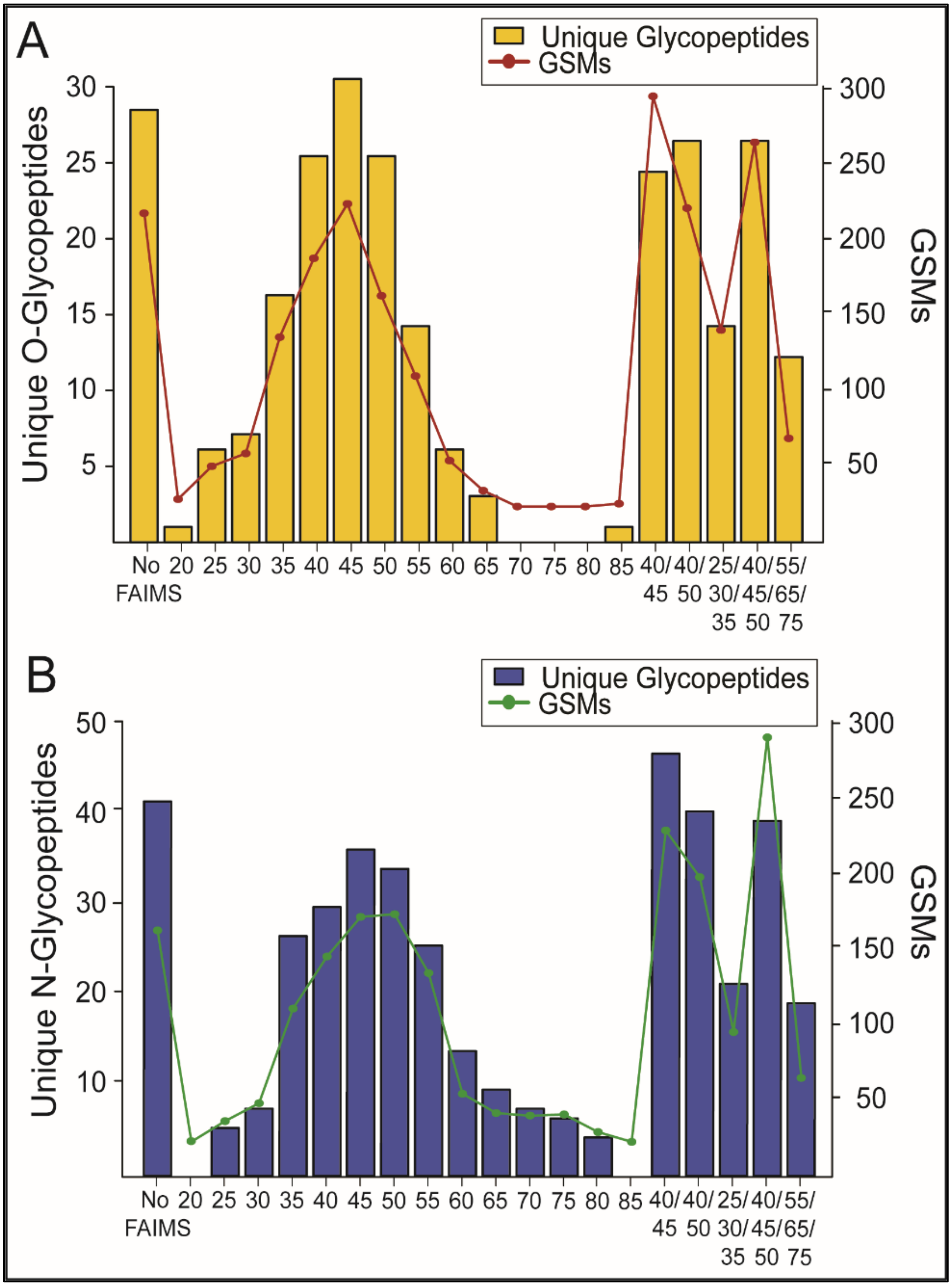


**Figure S9. N- and O-GSMs and UGPs identified in the glycoprotein mixture.** A mixture of six glycoproteins was generated, digested with trypsin, and subjected to FAIMS followed by MS analysis. Resulting RAW data was searched with Byonic, and the number of N- and O-GSMs scatterplot (O-, red; N-, green) and UGPs (bar graph; O-, yellow; N-, blue) was compared. N- and O-glycopeptide results are depicted in A) and B) respectively.

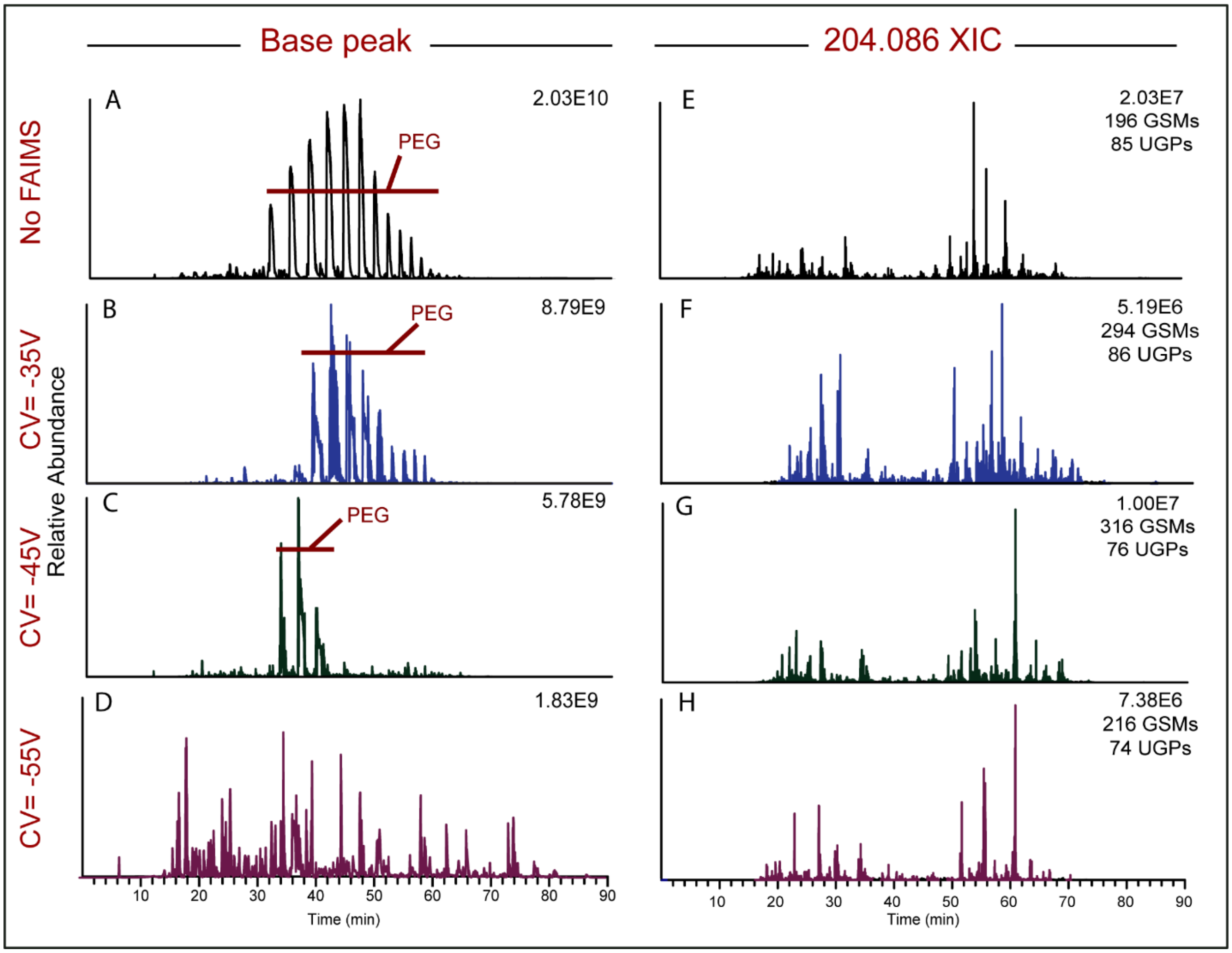


**Figure S10. Presence of PEG in the glycoprotein enrichment of the platelet sheddome across different CVs.** Experiments with and without FAIMS separation were performed using a lectin enriched platelet sheddome. Chromatograms were extracted using Thermo Scientific XCalibur Qual Browser. The left panel depicted the chromatographic base peak of experiments without FAIMS (A) and CVs -35V (B), -45V (C) and -55V (D). Normalized intensity is also depicted in the top right corner of each chromatogram. The right panel represented the extracted ion chromatogram of m/z 204.0867 (HexNAc oxonium ion) for control experiment (E) and CVs -35V (F), -45V (G) and -55V (H). Like the other panels, the normalized intensity is also pictured in the top right of each chromatogram, accompanied by the number of GSMs and UGPs identified for each setting.

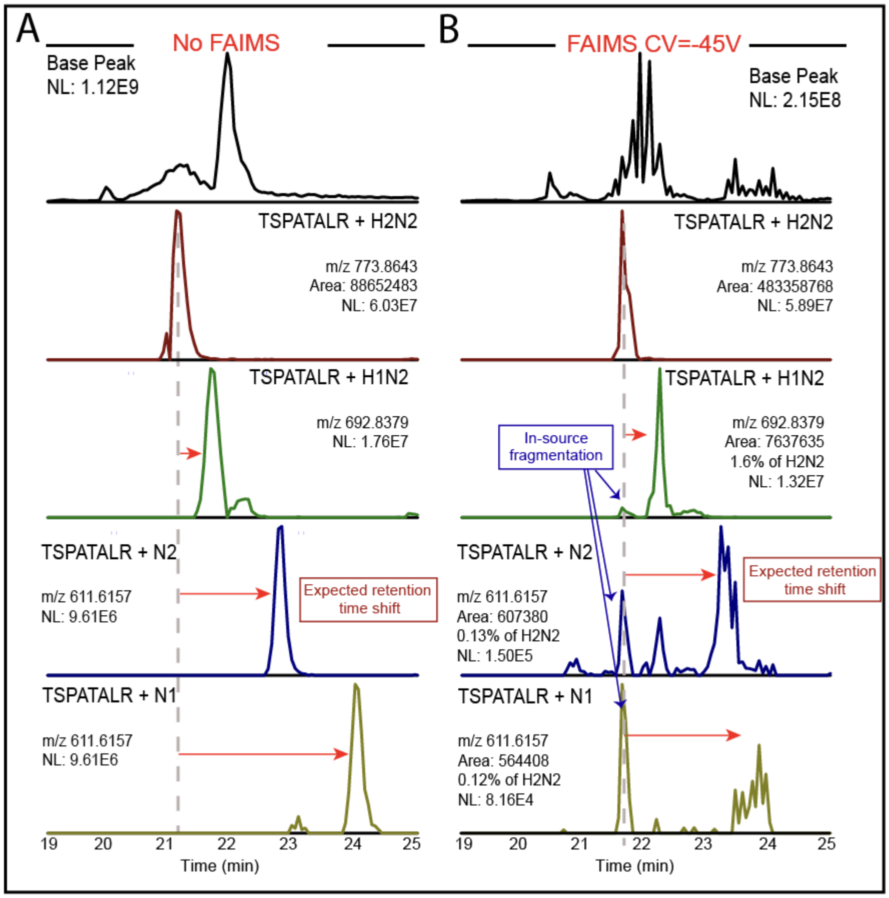


**Figure S11. Source fragmentation of O-glycopeptide TSPATALR with (IFF) and without FAIMS separation (ISF).** Recombinant podocalyxin was digested with mucinase SmE and trypsin, followed by manual analysis of XICs using Thermo Xcalibur Qual Browser.[2] Depicted here are the chromatographic base peak (black), extracted ion chromatogram of the doubly charged glycopeptide with glycan masses HexNAc2-Hex2 (red), HexNAc2-Hex1 (green), HexNAc2 (blue), and HexNAc2 (yellow). The percentages were calculated relative to that of the original precursor species (TSPATALR + HexNAc2-Hex2). Chromatograms depicted in (A) corresponded to the run without FAIMS separation, while chromatograms in (B) corresponded to setting -45V. Alongside each chromatogram, the glycopeptide and glycan composition, m/z, integrated area, and normalized abundance of peaks assigned as ISF/IFF are noted. A dashed gray line at 21 minutes indicated the retention time of the original precursor. Red arrows illustrated expected retention shift. Blue arrows indicated peaks originating from ISF/IFF.

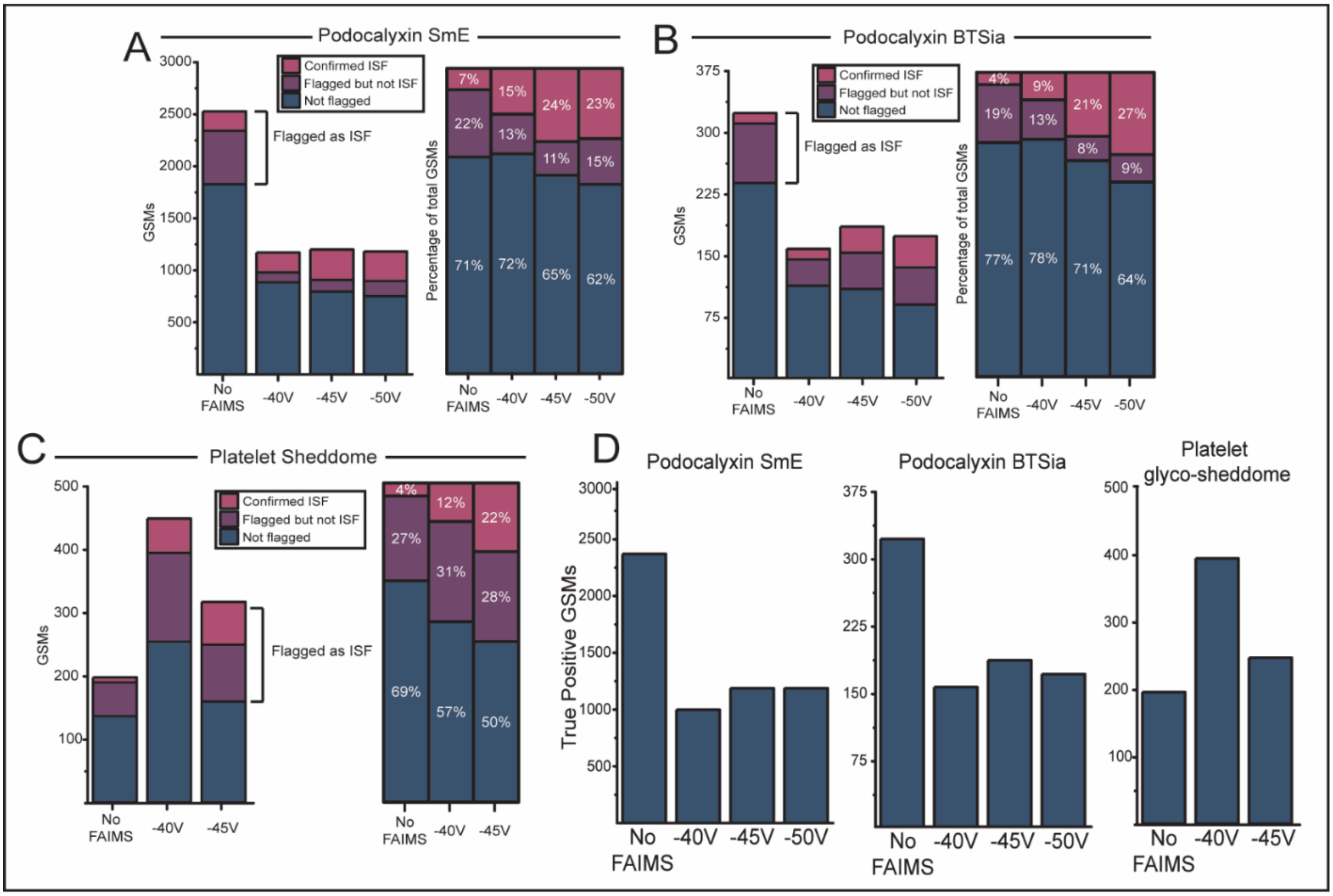


**Figure S12. ISF/IFF evaluation in three sample types.** Recombinant podocalyxin was digested with mucinase SmE[2] (A) and BTSia[4] (B), along with the lectin enriched glyco-sheddome of platelets[8] (C). Search results were analyzed with Glyco-SourceFragFinder, which flagged identifications that eluted within a minute of another identification with the same peptide backbone and larger glycan mass. Following this analysis, these plausible instances of ISF/IFF were validated by comparing their chromatographic profiles. GSMs that were not flagged by Glyco-SourceFragFinder are depicted in blue; GSMs flagged, but not confirmed as ISF/IFF, are labeled in purple; confirmed ISF/IFF is depicted in pink. The bar graph on the left (A-C) represented the counts in terms of total GSMs, while the graph on the right pictured the normalized percentage compared to the total number of GSMs. (D) True positive GSMs after subtracting the confirmed instances of ISF/IFF.

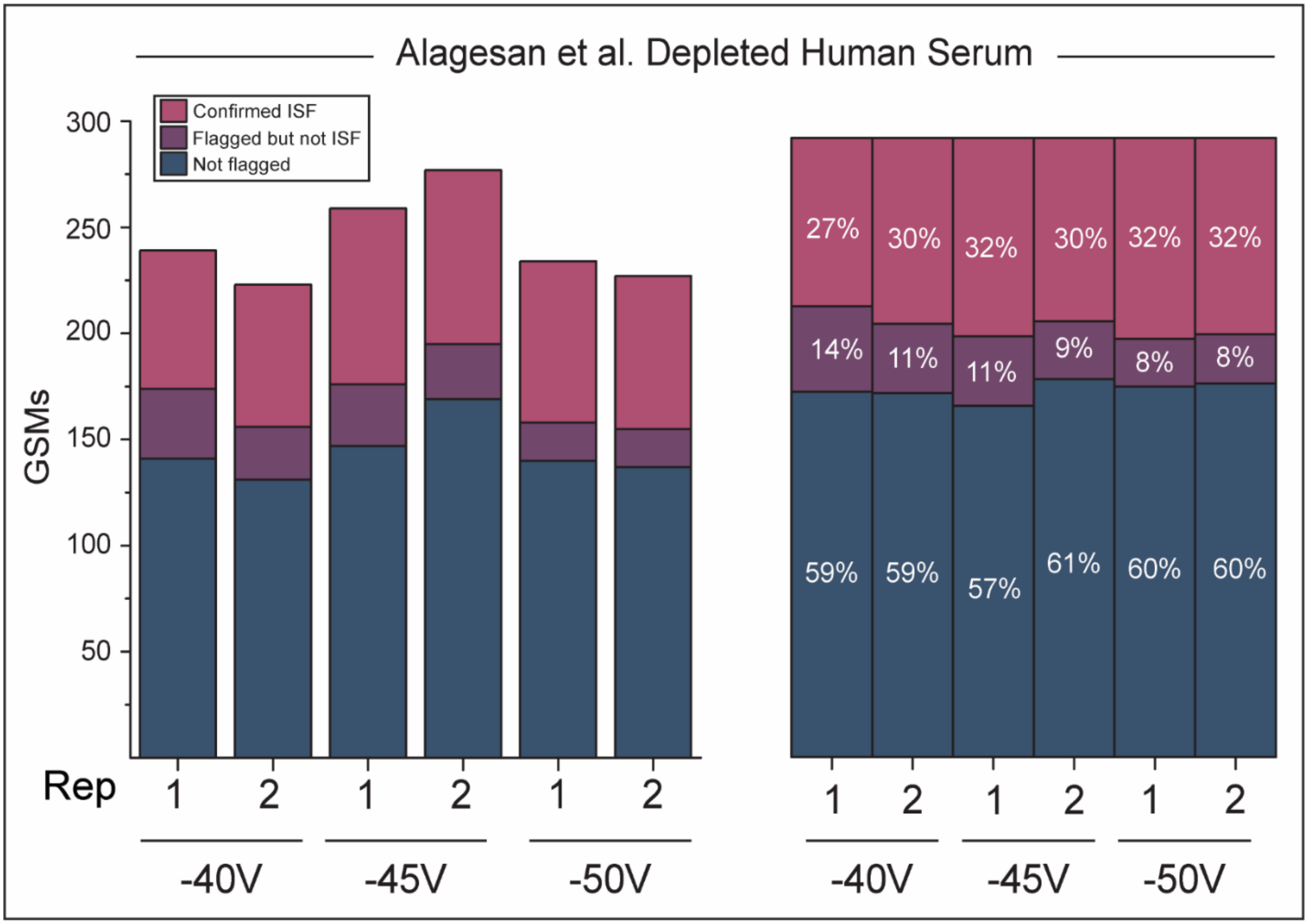


**Figure S13. Instances of IFF observed in the analysis of depleted human serum.** Depleted human serum RAW data was downloaded from the PRIDE repository with identifier PXD038673 and searched with Byonic[5] using the same settings applied for the platelet glyco-sheddome.[1] Search results were analyzed with Glyco-SourceFragFinder and plausible instances of IFF were validated. GSMs that were not flagged by Glyco-SourceFragFinder are depicted in blue; GSMs flagged, but not confirmed as IFF are labeled in purple; confirmed IFF is depicted in pink. The bar graph on the left represented the counts in terms of total GSMs, while the graph on the right portrayed the normalized percentages.

### References


(1) Alagesan, K.; Ahmed-Begrich, R.; Charpentier, E. *Improved N- and O-Glycopeptide Identification Using High-Field Asymmetric Waveform Ion Mobility Spectrometry (FAIMS)*; preprint; Biochemistry, 2022. https://doi.org/10.1101/2022.12.12.520086.

(2) Chongsaritsinsuk, J.; Steigmeyer, A. D.; Mahoney, K. E.; Rosenfeld, M. A.; Lucas, T. M.; Ince, D.; Kearns, F. L.; Battison, A. S.; Hollenhorst, M. A.; Shon, D. J.; Tiemeyer, K. H.; Attah, V.; Kwon, C.; Bertozzi, C. R.; Ferracane, M. J.; Amaro, R. E.; Malaker, S. A. Glycoproteomic Landscape and Structural Dynamics of TIM Family Immune Checkpoints Enabled by Mucinase SmE. bioRxiv February 3, 2023, p 2023.02.01.526488. https://doi.org/10.1101/2023.02.01.526488.

(3) Lu, L.; Riley, N. M.; Shortreed, M. R.; Bertozzi, C. R.; Smith, L. M. O-Pair Search with MetaMorpheus for O-Glycopeptide Characterization. *Nat Methods* **2020**, *17* (11), 1133–1138. https://doi.org/10.1038/s41592-020-00985-5.

(4) Shon, D. J.; Malaker, S. A.; Pedram, K.; Yang, E.; Krishnan, V.; Dorigo, O.; Bertozzi, C. R. An Enzymatic Toolkit for Selective Proteolysis, Detection, and Visualization of Mucin-Domain Glycoproteins. *Proc. Natl. Acad. Sci. U.S.A.* **2020**, *117* (35), 21299–21307. https://doi.org/10.1073/pnas.2012196117.

(5) Bern, M.; Kil, Y. J.; Becker, C. Byonic: Advanced Peptide and Protein Identification Software. *Curr Protoc Bioinformatics* **2012**, *CHAPTER*, Unit13.20. https://doi.org/10.1002/0471250953.bi1320s40.

(6) Ahmad Izaham, A. R.; Ang, C.-S.; Nie, S.; Bird, L. E.; Williamson, N. A.; Scott, N. E. What Are We Missing by Using Hydrophilic Enrichment? Improving Bacterial Glycoproteome Coverage Using Total Proteome and FAIMS Analyses. *J. Proteome Res.* **2021**, *20* (1), 599–612. https://doi.org/10.1021/acs.jproteome.0c00565.

(7) Khan, A.; Mathelier, A. Intervene: A Tool for Intersection and Visualization of Multiple Gene or Genomic Region Sets. *BMC Bioinformatics* **2017**, *18* (1), 287. https://doi.org/10.1186/s12859-017-1708-7.

(8) Hollenhorst, M. A.; Tiemeyer, K. H.; Mahoney, K. E.; Aoki, K.; Ishihara, M.; Lowery, S. C.; Rangel-Angarita, V.; Bertozzi, C. R.; Malaker, S. A. Comprehensive Analysis of Platelet Glycoprotein Ibα Ectodomain Glycosylation. *Journal of Thrombosis and Haemostasis* **2023**. https://doi.org/10.1016/j.jtha.2023.01.009.
